# EVALUATION OF ANTIMICROBIAL AND ANTIPROLIFERATIVE ACTIVITIES OF ACTINOBACTERIA ISOLATED FROM THE SALINE LAGOONS OF NORTHWEST PERU

**DOI:** 10.1101/2020.10.07.329441

**Authors:** Rene Flores Clavo, Nataly Ruiz Quiñones, Álvaro Tasca Hernandez, Ana Lucia Tasca Gois Ruiz, Lucia Elaine de Oliveira Braga, Zhandra Lizeth Arce Gil, Luis Miguel Serquen Lopez, Jonas Henrique Costa, Taícia Pacheco Fill, Marcos José Salvador, Fabiana Fantinatti Garboggini

## Abstract

The unexplored saline lagoons of the north of Peru harbor a rich microbiome, due to reported studies of different extreme environments around the world. In these regions, there are several ecosystems and microhabitats not yet explored, and little is known about the diversity of actinobacteria and other microorganisms. We suggest that the endemic bacteria present in this extreme environment could be source of active molecules with anticancer, antimicrobial, antiparasitic properties. Using phenotypic and genotypic characterization techniques including the 16S rRNA were identified into the genera *Streptomyces* 39 (78%), *Pseudonocardia* 3 (6%), *Staphylococcus* 4 (8%), *Bacillus* 2 (4%), and *Pseudomonas* 2 (4%). All isolated bacteria for the genotypic data were preliminarily identified. Actinobacteria strains were found dominantly in both sites (Lagoon1-3 = 16 isolates and lagoon 4 = 12 isolates). Phylogenetic analysis revealed that 28 isolates were exclusively affiliated to eleven different clusters of Actinobacteria of the major genus *Streptomyces*. Three *Streptomyces* sp. strains M-92, B-146, and B-81, were tested for antibacterial and antiproliferative activities. The results showed antiproliferative activities against three tumor cell lines, U251 glioma; MCF7 breast; NCI-H460 lung non-small type of cells, and the antibacterial activity to *Staphylococcus aureus* ATCC 6538, *E. coli* ATCC 10536, and *Acinetobacter baumanni* AC-972 which is resistant to multiple drugs. The promising results belong to *Streptomyces* sp. B-81 strain in the R2A medium using a doxorubicin with control positive, the best result was from the latter (TGI = 0,57 µg/mL) for glioma; NCI-H460 lung of type non-small cells (TGI = 0,61 µg/mL), and breast cancer (TGI =0,80 µg/mL), this strain was selected to be fractionated because it had better antiproliferative and antibacterial activity, and its fractions were evaluated concerning antiproliferative activity against nine types of tumor cells and one non-tumor. The methanolic fraction showed a better result in the antiproliferative activity and was able to inhibit U251 (glioma) (TGI = 38.3 µg/mL), OVCAR-03 (ovary) (TGI = 62.1 µg/mL), and K562 (leukemia) (TGI = 81.5 µg/mL). The methanol 50% - acetate 50% fraction (Fraction 4) inhibited U251 (glioma) (TGI = 73.5 µg/mL) and UACC-62 (melanoma) (TGI = 89.4 µg/mL). Moreover, the UHPLC-MS/MS data and molecular networking of *Streptomyces sp*. B-81 isolate extract revealed the production cholic acid, Lobophorin A, Lobophorin B, Lobophorin E, Lobophorin K and compound 6. Extremophilic environments such as the Mórrope and Bayovar Salt Flats are promising sources of new bacteria with promising pharmaceutical potential; These compounds could be useful to treat various infectious diseases or even some type of cancer.

## 1. Introduction

Perú has extreme environments such as the salt marshes located on the coast, center, and south of the country. In these regions, there are several ecosystems and microhabitats, whose studies are still restricted and report the diversity of bacteria and other organisms (Caceres and Legendre, 2009; Flores and Pisfil, 2014; Leon et al. 2016). Among other microorganisms, the members of the phylum Actinobacteria can be found in all kinds of extreme environments. *Micromonospora, Actinomadura*, and *Nocardiopsis* were reported from saline soils of the ephemeral salty lakes in Buryatiya (Lubsanova et al. 2014). *Streptomyces, Nocardiopsis*, and *Nocardioides* were isolated from the Regions Western Ghats of India (Siddhart et al. 2020). Concurrent to this, there is still little information documented on isolated substances and their biological activities of the microorganisms isolated from saline environments. *Micromonospora, Streptomyces, Salinispora*, and *Dietzia* were isolated from the coastal zone of the geographically remote young volcanic Easter Island Chile. Besides, actinobacterial halophilic and halotolerant strains show heterogeneous physiological characteristics among different genera because these bacteria have the ability to synthesize secondary metabolites to cope with conditions of high salinity and the different temperatures present in their environments (Binayke et al. 2018; Oueriaghli et al. 2018; Boyadzhieva et al. 2018). These extreme conditions favor the development of metabolically competitiveness to the production enzymes, which can help the bacterial population to adapt to high salinity (Kim et al. 2019). Furthermore, these microorganisms can also perform essential processes such as carbon cycle, metal transfer, and the removal of organic pollutants to higher trophic levels (Fiedler et al. 2005; Blunt et al. 2007). In addition, this microbial group presents a unique ability to produce new products, mainly antibiotics (Berdy, 2005; Strohl, 2004).

The search for these microorganisms has been mainly related to the production of antibiotics and antitumor substances (Newman and Cragg, 2007). There are about 22.500 biologically active substances obtained from microorganisms, 45% of which are represented by actinomycetes, with *Streptomyces* being the producer of 70% of them (Hayakawa et al. 2007 and Valliappan et al. 2014). According to Lam (2006), there are new actinobacteria from habitats not yet explored or little explored to be the source of new bioactive secondary metabolites.

The bioguided study strategy helps us find a wide variety of microorganisms that have biotechnological potential (Maldonado et al. 2005). This study reports the isolation and phylogenetic analysis of a collection of bacterial isolates from the saline lagoons from the Northern Peru, and their potential as producers of secondary metabolites with antimicrobial and antiproliferative activities.

## 2. Materials and methods

### 2.1 Site description and sampling

Sample collection at Mórrope saline lagoons was carried out in December 2012 the zone “1”, in January 2013 the zone “2”, and in July 2014 the zone “3”, while samples were collected at Bayovar saline lagoons in March 2015 the zone “4” (Figure 1). The present work did not need permits from a competent authority, since it does not include any protected species, because the standard is currently in the process of being implemented to achieve its registration, likewise the sampling areas are lagoons that due to their nature are in the process of adjudication to the competent governmental organism through the Peruvian Ministry of the Environment.

**Fig 1.**
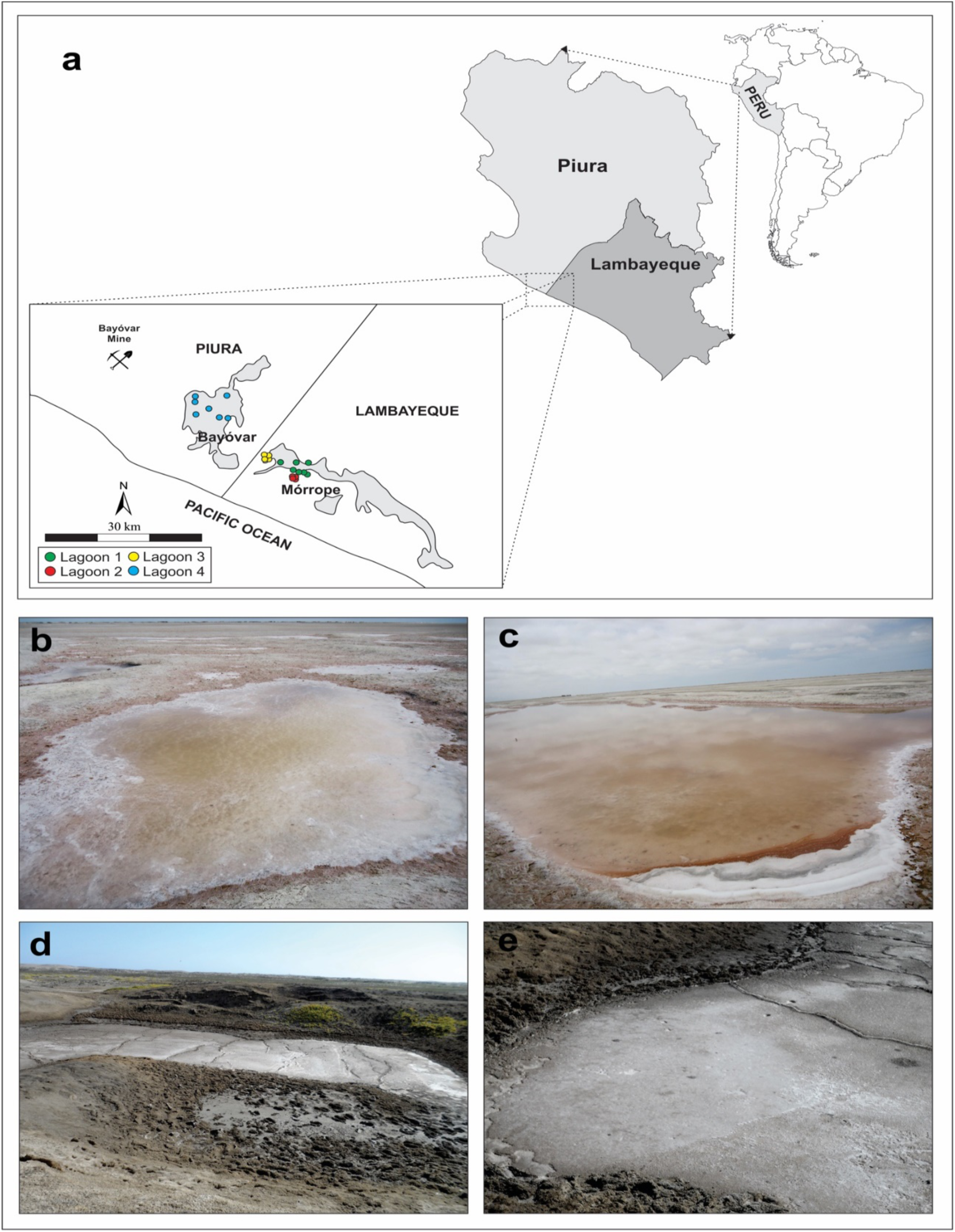
a = Geographic location of sampling sites; b and c = lagoons 1, 2 and 3 (State Mórrope); and e = lagoon 4 (State Bayovar).

The sources and places from which samples were obtained are detailed in Table 1. Samples were collected aseptically, placed in sterile plastic bags, and kept refrigerated at 4 °C for further processing in the Laboratory of Bacteriology the Regional Hospital of Lambayeque, a flowchart depicting the methodological strategy adopted in this work is shown in Figure 2.

**Table 1.** Data related to saline lagoons samples.

| **Mórrope lagoons** |  |  |  |  |
| --- | --- | --- | --- | --- |
| **Collection zone** | **Sample type** | **Geographic location** |  |  |
| **Zone 1**<br>**(December 2012)** | A; AES | 6°10S;80°35W<br>(3 samples x triplicate) | 6°11S;80°37W<br>(3 samples x triplicate) | 6°9S;80°39W<br>(3 samples x triplicate) |
| **Zone 2**<br>**(January 2013)** | AES; SE | 6°19S;80°28W<br>(3 samples x triplicate) | 6°17S;80°26W<br>(3 samples x triplicate) | 6°18S;80°27W<br>(3 samples x triplicate) |
| **Zone 3**<br>**(December 2014)** | A; SE | 6°08S;80°50W<br>(3 samples x triplicate) | 6°08S;80°51W<br>(3 samples x triplicate) | 6°08S;80°40W<br>(3 samples x triplicate) |
| **Bayovar lagoons** |  |  |  |  |
| **Zone 4**<br>**(March 2015)** | A; AES; SE | 6°23S;80°66W<br>(3 samples x triplicate) | 6°25S;80°26W<br>(3 samples x triplicate) | 6°23S;80°78W<br>(3 samples x triplicate) |

A= (saline water), AES= (water and sediment), SE= (sediment).

**Fig. 2.**
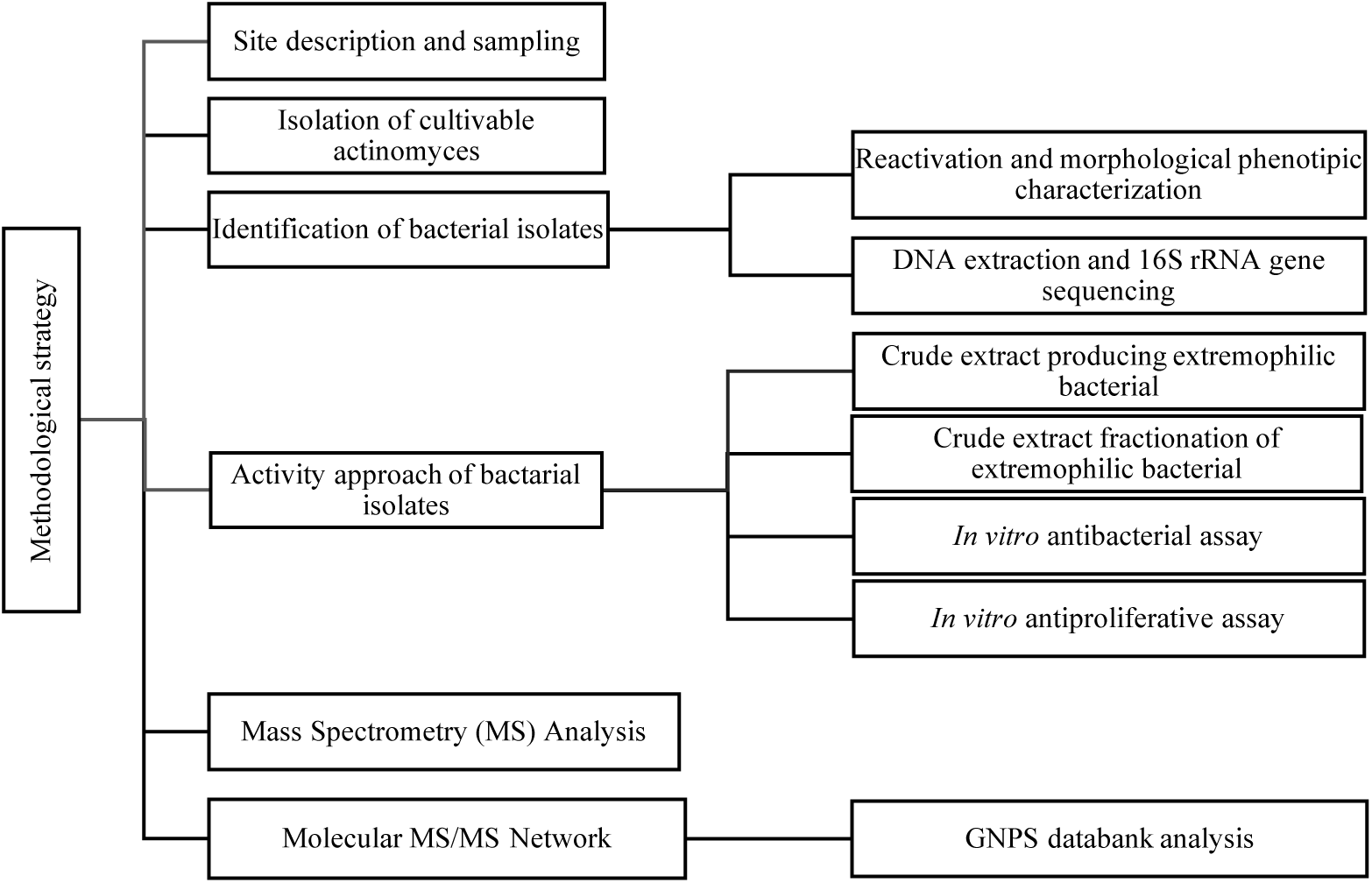
Flowchart depicting the methodological strategy adopted in this study.

### 2.2 Isolation of cultivable actinomyces

An aliquot of 10.0 mL of saline lagoons sample was transferred to an Erlenmeyer containing 10.0 mL of salt broth (0.6% yeast extract, 2% glucose, 5% peptone, 3% meat extract supplemented with chloramphenicol and 1% fluconazole at pH 7.0). All Erlenmeyer were homogenized and maintained at 50 °C in a water bath for 60 min to reduce the pollutant load (Pisano et al. 1989 and Takizawa et al. 1993). Then, they were incubated at 28 °C for 7 to 30 days under aerobic conditions; after, the microorganisms were isolated in the media: modified saline agar (1.0 g yeast extract, 5.0 g peptone, and 15.0 g of agar at pH 7.6 ± 0.2) and trypticase soy agar (Bacto™) supplemented with saline water. The isolates were preserved in 20% glycerol and kept at -80 °C cryopreserved. The isolates were reactivated and stored in a refrigerated chamber at -20 ° C to the transport to the Microbial Resources Division of the Pluridisciplinary Center for Chemical, Biological and Agricultural Research (CPQBA) in August 2015.

### 2.3 Identification of bacterial isolates

#### 2.3.1 Reactivation and morphological phenotypic characterization of bacterial strains

The reactivation of the isolates was carried out in R2A broth (Difco ref. 234000), supplemented with artificial seawater (ASW) NaCl 5% (0.1 g potassium bromide, 23.48 g sodium chloride, 10.61 g of magnesium chloride, 1.47 g of calcium chloride, 0.66 g of potassium chloride, 0.04 g of strontium chloride, 3.92 g of sodium sulfate, 0.19 g of sodium bicarbonate, 0.03 g of boric acid all in 1 L of distilled water) at 28 °C for 28 days, the colonies were observed under a microscope and stereoscope for macro morphological characterization and after Gram staining, the cells were observed under an optical microscope. The bacteria were preserved in 20% glycerol and kept at -80 °C.

#### 2.3.2 DNA extraction and 16S rRNA gene sequencing

An isolated colony was used for extraction of genomic DNA, according to the method described by Pospiech and Neuman, 1995, with some modifications. Amplification of the 16S ribosomal RNA gene was done by means of a polymerase chain reaction (PCR) using the pair of primers p10f (5’ GAG TTT GAT CCT GGC TCA G 3’) and p1401r (5’ CGG TGT GTA CAA GGC CCG GGA ACG 3’) or p1492r (5’-ACC TTG TTA CGA CTT 3’) (Lane et al. 1985) homologous to conserved regions of the bacterial 16S ribosomal RNA gene. Conditions used in the amplification reaction were: 2.5 µL of 10X buffer (without Mg^+2^), 0.75 µL of MgCl_2_ (2.5 mM), 0.2 µL dNTP (2.5 mM each), 0.5 µL of each primer (20 µM), 0.2 µL of Taq DNA polymerase (Invitrogen) and 5.0 µL of the DNA template (50-100 ng/ µL) and deionized and sterilized water by filling the volume to 25 µL of reaction. The PCR amplification was an initial denaturation step of 5 min at 95°C, followed by 30 cycles of 1 min at 94°C for denaturation, 1 min at 55°C for annealing, 3 min at 72°C for extension and 3 min at 72°C for final extension in an Eppendorf thermal cycler. The obtained products were analyzed and visualized in agarose gel electrophoresis, purified with mini columns (GFX PCR DNA & gel band purification kit, GE Healthcare), and sequenced in ABI3500XL Series automatic sequencer (Applied Biosystems). Sequencing reactions were performed with the Big Dye Terminator Cycle Sequencing Ready Reaction Kit (Applied Biosystems) according to the manufacturer’s specifications.

Partial sequences of the 16S ribosomal RNA gene obtained from each isolate were assembled into a contig and then compared to the sequences of organisms represented in EZBioCloud 16S Database (https://www.ezbiocloud.net/) using the “Identify” service (Yoon et al. 2017), and species assignment were based on closest hits (Jeon et al. 2014). 16S ribosomal RNA gene sequences retrieved from the database and related to the unknown organism gene were selected for alignment in the Clustal X program (Thompson et al. 1997), and phylogenetic analyzes were performed using the Mega version 6.0 program (Tamura et al. 2013). The evolutionary distance matrix was calculated with the model of Kimura-2 parameters (Kimura, 1980), and the phylogenetic tree constructed from the evolutionary distances calculated by the Neighbor-Joining method (Saitou and Nei, 1987), with bootstrap values from 1000 resampling.

### 2.4 Activity approach of bacterial isolates

#### 2.4.1 Crude extract producing extremophilic bacterial

Secondary metabolites of bacterial isolates were extracted with ethyl acetate. A pre-inoculum of the bacteria was performed in 5.0 mL of R2A Broth medium (Himedia ref. 1687) with AWS NaCl 5% and incubated at 28 °C for 5 to 7 days. After the growth of the culture, the total volume was transferred to an Erlenmeyer containing 500.0 mL of the same culture medium and incubated for 7 days under agitation at 150 rpm, followed by 23 days without agitation. After this period, 500.0 mL of ethyl acetate were added to the medium containing the bacterial growth and ruptured in ultra turrax basic (IKA ref. 02H2063.08.CC) at 7,000 rpm for 15 min. The organic fraction was obtained and the crude extract was concentrated in the rotary evaporator (R-215 Buchi), under vacuum and temperature of 40 °C, until the solvent was completely dried and stored at 4°C. The crude extracts of three strains representatives were taken for antimicrobial and antiproliferative activity tests. Simultaneously, the crude extracts also were analyzed by liquid chromatography coupled to the mass spectrometer (Versamax, molecular manufacturer Devices).

#### 2.4.2 Crude extract fractionation of the extremophilic bacterial

The crude extracts of the isolates were fractionated using a vacuum chromatographic column with C18. The initial crude extract was solubilized in methanol. The mobile phases used were fraction 1 water (H_2_O); fraction 2 water: methanol (H_2_O: MeOH 1:1 v/v); fraction 3 methanol (MeOH); fraction 4 MeOH: ethylacetate (EtOAc) (1:1 v/v); fraction 5 EtOAc 100%; fraction 6 EtOAc: Glacial acetic acid 1%. These fractions were dried in rotary evaporator at 45 °C, under vacuum weighted and used for activity antibacterial and antiproliferative tests.

#### 2.4.3 *In vitro* antibacterial activity assay

Three crude bacterial extracts (M-92, B-81 and B-146) were tested as antimicrobial producers using the minimum inhibitory concentration (MIC) assay, following the protocol reported by Siddharth and Rai 2018. The crude extracts partially diluted in 1% Dimethylsulfoxide (DMSO) 1 mg/mL – 3.9 µg/mL, and sterile broth were added into pre-coated microbial cultures, completed a total volume of 200 µL. The plate was incubated at 37 °C and room temperature; the lowest concentration of extract, which completely inhibited the bacterial growth was considered as MIC. Each biological assay was performed in triplicate. The pathogenic bacteria used in this test were *Escherichia coli* ATCC 10536, *Staphylococcus aureus* ATCC 6538, and *Acinetobacter baumannii* with code AC-972, the strain was not acquired prospectively, the source of origin comes from the bank of MDRs isolated from a patient with pneumonia from the UCI of the Hospital Regional Lambayeque, details such as patient data linked to the sample were anonymized in order to obtain access

#### 2.4.4 *In vitro* antiproliferative activity assay

This assay aimed to detect anticancer activities by evaluating antiproliferative action in human tumor cells (Monks et al. 1991). *In vitro* tests with the crude extract of *Streptomyces* sp. M-92, B-81, and B-146 on human tumor cell lines of different origins and a non-tumor cell line. Human tumor cell lines U251 (glioma), UACC-62 (melanoma), MCF-7 (breast), NCI-ADR/RES (ovarian expressing phenotype with multiple drugs resistance), 786-0 (renal), NCI-H460 (lung, non-small cells), PC-3 (prostate), OVCAR-03 (ovarian), K562 (leukemia) and a non-tumor cell line HaCaT (keratinocyte) were obtained from the National Cancer Institute Frederick, Molecules (MD, USA). Stock cultures were grown in medium containing 5 mL of RPMI 1640 (GIBCO BRL, Gaithers-Burg, MD, USA) supplemented with 5 % fetal bovine serum (FBS, GIBCO) at 37 °C with 5% CO_2_. Penicillin: streptomycin (1000 µg·L−1:1000 U·L−1, 1 mL·L−1) was added to the experimental cultures. Cells in 96-well plates (100 µL cells well−1) were exposed to the extracts in DMSO (Sigma-Aldrich) /RPMI (0.25, 2.5, 25, and 250 µg·mL−1) at 37 °C, and 5% CO_2_ in the air for 48 h. The final DMSO concentration (0.2% in higher concentration) did not affect cell viability. Before (T_0_) and after (T_1_) sample application, cells were fixed with 50% trichloroacetic acid (Merck), and cell proliferation was determined by the spectrophotometric quantification (540 nm) of cellular protein content using sulforhodamine B assay. Using the concentration-response curve for each cell line, the values of the sample concentration required to produce total growth inhibition or cytostatic effect (TGI) were determined through non-linear regression analysis using software ORIGIN 8.6® (OriginLab Corporation, Northampton, MA, USA).

### 2.5 Mass Spectrometry analysis

*Streptomyces sp*. B-81 extract was resuspended in 1 mL of methanol (HPLC grade) and 100 µL were diluted in 900 µL of methanol. The final solution was filtered through 0.22 µm into vials. Ultra-high pressure liquid chromatography-mass spectrometry (UHPLC-MS) analyses were performed in a Thermo Scientific QExactive® Hybrid Quadrupole-Orbitrap Mass Spectrometer. Parameters: positive mode, capillary voltage at +3.5 kV; capillary temperature at 250 °C; S-lens of 50 V and *m/z* range of 133.40-2000.00. Tandem mass spectrometry (MS/MS) was performed using normalized collision energy (NCE) of 30 eV and 5 precursors per cycle were selected. Stationary phase: Thermo Scientific Accucore C18 2.6 µm (2.1 mm x 100 mm) column. Mobile phase: 0.1% formic acid (A) and acetonitrile (B). Eluent profile (A: B) 0-10 min, gradient from 95:5 up to 2:98; held for 5 min; 15-16.2 min gradient up to 95:5; held for 8.8 min. Flow rate: 0.2 mL min^-1^. The injection volume of the samples was 3 µL. Operation and spectra analyses were conducted using Xcalibur software (version 3.0.63) developed by Thermo Fisher Scientific.

### 2.6 Molecular MS/MS network

A molecular network for *Streptomyces sp*. B-81 was created using the online workflow at GNPS (https://gnps.ucsd.edu/). The data were filtered by removing all MS/MS peaks within ± 17 Da of the precursor *m/z*. MS/MS spectra were window filtered by choosing only the top 6 peaks in the ± 50 Da window throughout the spectrum. The data was then clustered with MS-Cluster with a parent mass tolerance of 0.02 Da and a MS/MS fragment ion tolerance of 0.02 Da to create consensus spectra. A network was then created where edges were filtered to have a cosine score above 0.5 and more than 5 matched peaks. Further edges between two nodes were kept in the network only if each of the nodes appeared in each other’s respective top 10 most similar nodes. The spectra in the network were then searched against GNPS’ spectral libraries. The library spectra were filtered in the same manner as the input data. All matches kept between network spectra and library spectra were required to have a score above 0.5 and at least 5 matched peaks (Wang et al. 2016).

## 3. Results and discussion

### 3.1 Isolation, identification and selective of bacterial species from the northern saline lagoons of Perú

In total, 166 pure cultures showing different colony morphologies were obtained grown on R2A medium from saline water (A), water and sediment (AES), and sediment (SE). The morphological characteristics such as aerial mycelium, the morphological spore mass color, pigmentation of vegetative or substrate mycelium, and the production of diffusible pigment were used to classify 42 filamentous and 8 non filamentous to the genera *Streptomyces* 39 (78%), *Pseudonocardia* 3 (6%), *Staphylococcus* 4 (8%), *Bacillus* 2 (4%), and *Pseudomonas* 2 (4%) in Table 2, (Flores, 2017).

**Table 2.**
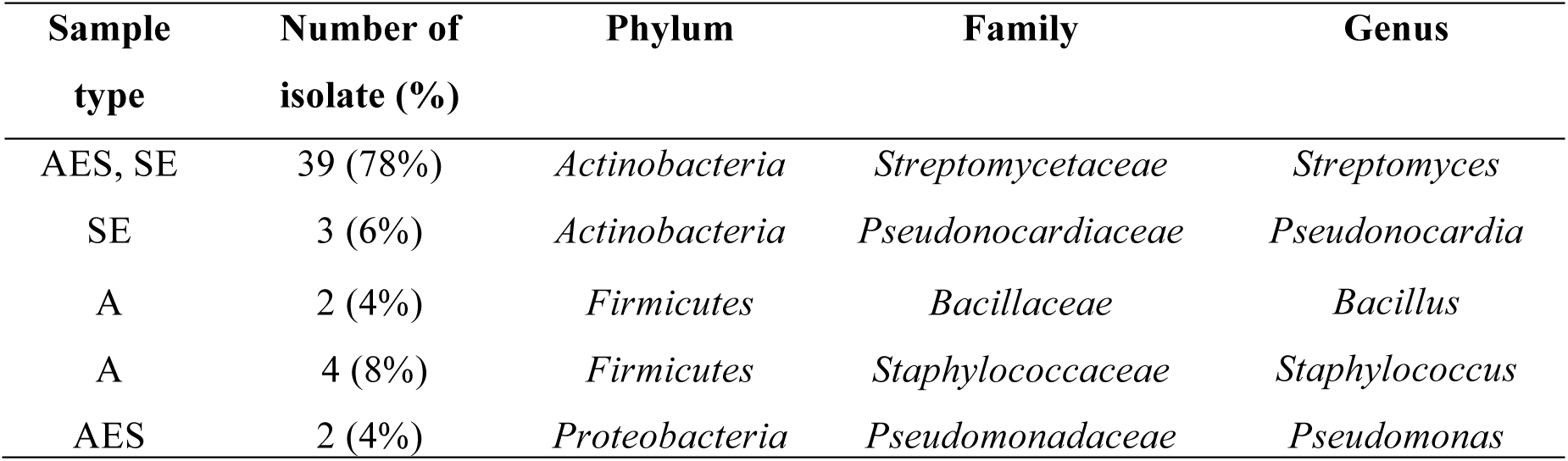
16S rRNA gene-based identification of bacteria isolate from lagoons Morrope and Bayovar.

The isolates were regrouped into 28 isolates due to their similar phenotypic characteristics and were identified based on the sequencing and alignment, from the 16S rRNA gene analyzes. They belong to the genus *Streptomyces* 27 isolates and 1 *Pseudonocardia* isolate, and 23 isolates showed a 16S rRNA gene sequence similarity below the 98.7% threshold for proposing as novel species (Table 3).

**Table 3.**
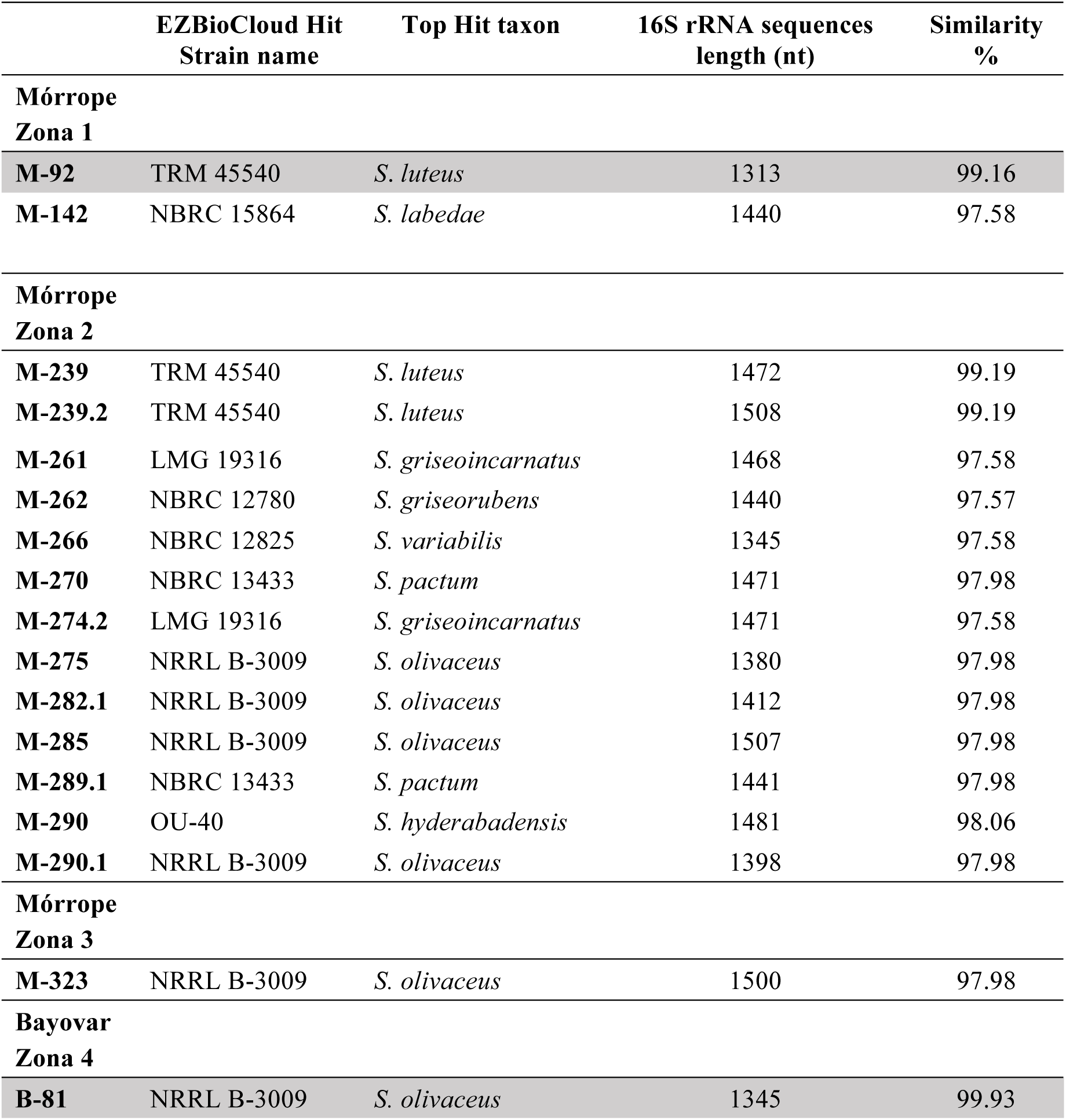
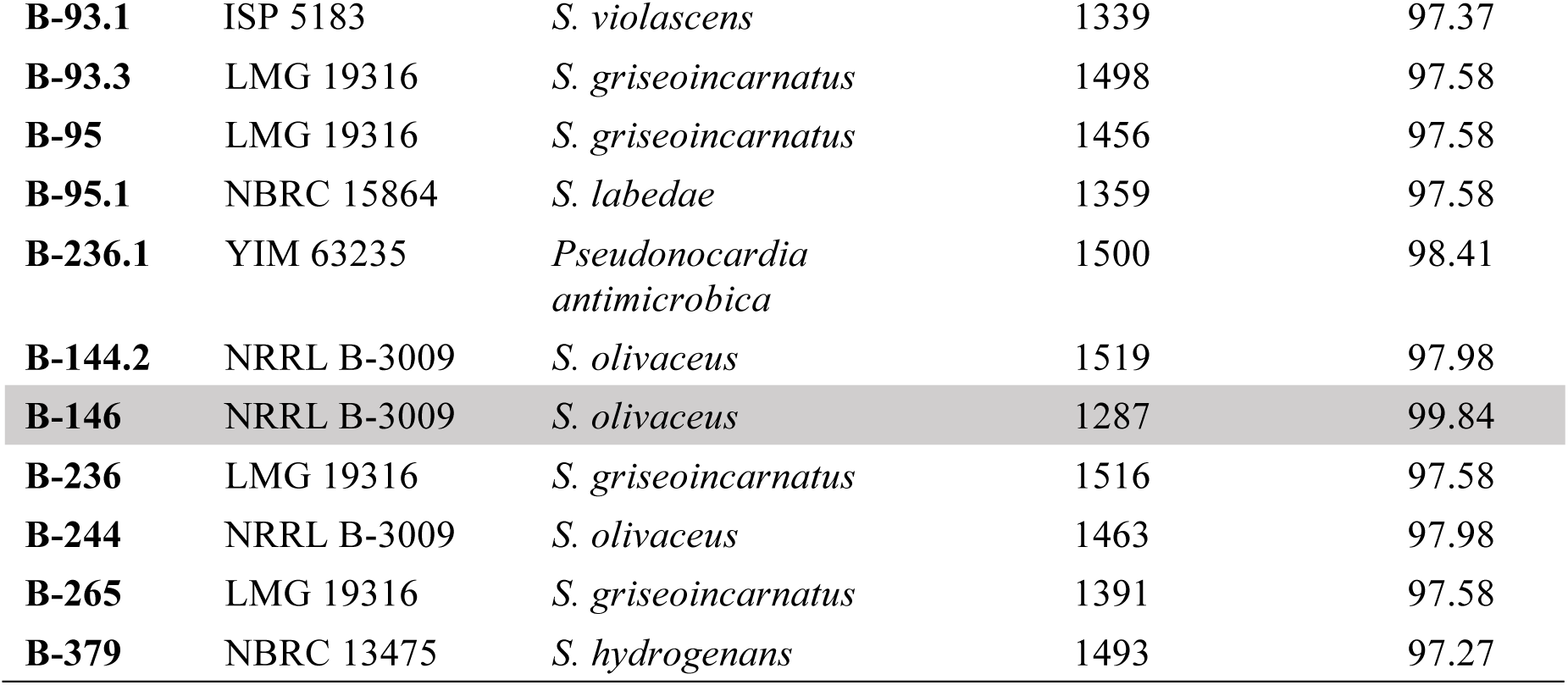
**Strain Actinobacterial isolate from different sample Bayovar and Mórrope saline lagoons, indicating the top hit taxon, sequences length nucleotides, and similarity percent of the isolates the “Identify” service best-hit classifier in the EZBioCloud 16S Database.**

|  | EZBioCloud Hit<br>Strain name | Top Hit taxon | 16S rRNA sequences<br>length (nt) | Similarity<br>% |
| --- | --- | --- | --- | --- |
| **Mórrope**<br>**Zona 1** |  |  |  |  |
| **M-92** | TRM 45540 | *S. luteus* | 1313 | 99.16 |
| **M-142** | NBRC 15864 | *S. labedae* | 1440 | 97.58 |
| **Mórrope**<br>**Zona 2** |  |  |  |  |
| **M-239** | TRM 45540 | *S. luteus* | 1472 | 99.19 |
| **M-239.2** | TRM 45540 | *S. luteus* | 1508 | 99.19 |
| **M-261** | LMG 19316 | *S. griseoincarnatus* | 1468 | 97.58 |
| **M-262** | NBRC 12780 | *S. griseorubens* | 1440 | 97.57 |
| **M-266** | NBRC 12825 | *S. variabilis* | 1345 | 97.58 |
| **M-270** | NBRC 13433 | *S. pactum* | 1471 | 97.98 |
| **M-274.2** | LMG 19316 | *S. griseoincarnatus* | 1471 | 97.58 |
| **M-275** | NRRL B-3009 | *S. olivaceus* | 1380 | 97.98 |
| **M-282.1** | NRRL B-3009 | *S. olivaceus* | 1412 | 97.98 |
| **M-285** | NRRL B-3009 | *S. olivaceus* | 1507 | 97.98 |
| **M-289.1** | NBRC 13433 | *S. pactum* | 1441 | 97.98 |
| **M-290** | OU-40 | *S. hyderabadensis* | 1481 | 98.06 |
| **M-290.1** | NRRL B-3009 | *S. olivaceus* | 1398 | 97.98 |
| **Mórrope**<br>**Zona 3** |  |  |  |  |
| **M-323** | NRRL B-3009 | *S. olivaceus* | 1500 | 97.98 |
| **Bayovar**<br>**Zona 4** |  |  |  |  |
| **B-81** | NRRL B-3009 | *S. olivaceus* | 1345 | 99.93 |
| **B-93.1** | ISP 5183 | *S. violascens* | 1339 | 97.37 |
| **B-93.3** | LMG 19316 | *S. griseoincarnatus* | 1498 | 97.58 |
| **B-95** | LMG 19316 | *S. griseoincarnatus* | 1456 | 97.58 |
| **B-95.1** | NBRC 15864 | *S. labedae* | 1359 | 97.58 |
| **B-236.1** | YIM 63235 | *Pseudonocardia antimicrobica* | 1500 | 98.41 |
| **B-144.2** | NRRL B-3009 | *S. olivaceus* | 1519 | 97.98 |
| **B-146** | NRRL B-3009 | *S. olivaceus* | 1287 | 99.84 |
| **B-236** | LMG 19316 | *S. griseoincarnatus* | 1516 | 97.58 |
| **B-244** | NRRL B-3009 | *S. olivaceus* | 1463 | 97.98 |
| **B-265** | LMG 19316 | *S. griseoincarnatus* | 1391 | 97.58 |
| **B-379** | NBRC 13475 | *S. hydrogenans* | 1493 | 97.27 |

Therefore, these strains should be designed for further taxonomic and analytical chemistry analyses to confirm their novelty at species rank and as a source of novel chemical entities. Two isolates were identified in lagoon 1 of the Morrope salt flats, and according to the EZBioCloud the isolate M-92 presented similarity with *Streptomyces luteus* TRM 45540 (99.16%), while the isolate M-142 presented similarity with *Streptomyces labedae* NBRC 15864 (97.58%). In lagoon 2 of the Morrope salt flats, 13 isolates were identified, of which M-239 and M-239.2 were similar to *S. luteus* TRM 45540 (99.19%), the isolates M-261 and M-274.2 presented similar to *S. griseoincarnatus* LMG 19316 (97.58%), the isolate M-262 showed similarity to *S. griseorubens* NBRC 12780 (97.57%), the isolate M-266 was similar to *S. variabilis* NBRC 12825 (97.58%), the isolates M-270 and M-289.1 were similar to *S. pactum* NBRC 13433 (97.98%), the isolate M-290 presented similar to *S. hyderabadensis* OU-40 (98.06%), and finally four isolates, M-275, M-282.2, M-285 and M-290.1 were similar to *S. olivaceus* NRRL B-3009 (97.98%). From the lagoon 3, also belongs to the Morrope salt flats, it was identified one isolate M-323 similar to *S. olivaceus* NRRL B-3009 (97.98 %). Finally, from lagoon 4, belongs to the Bayovar salt flats, 12 isolates were identified, four isolates were similar to *S. olivaceus* NRRL B-3009, B-81 (99.93%0, B-144.2 (97.98%), B-146 (99.84%), and B-244 (97.98%). Also, in the lagoon 4 the isolate B-93, was similar to *S. violascens* ISP 5183 (97.37%), and other four isolates presented similar *S. griseoincarnatus* LMG 19316, B-93.3, B-95, B-236, B-265 (97.58%), the isolate B-95.1 was similar to *S. labedae* NBRC 15864 (97.58%), and the isolate B-379 was similar to *S. hydrogenans* NBRC 13475 (97.27%). Besides Streptomyces, another of the actinobacteria genus was recovered to the Bayovar salt flats. The isolate B-236.1 was similar to *Pseudonocardia antimicrobica* YIM 63235 (97.27%) and our work is the first to report this genus present in the Bayovar salt flats (Table 3).

In the Morrope salt flats, eight different groups of *Streptomyces* were identified, whereas in the Bayovar saline only five groups of *Streptomyces* were recovered, however, a different genus like *Pseudonocardia*, whichhas been reported by Zhang et al. (2016) as producers of γ-Butyrolactones that previously had only been reported for Streptomyces genera and in plants was recovered in this study (Table 3). Our culture-based approach using a pre-enrichment to decrease the bacterial load agrees with the reports described in the literature, in which the isolation of halophilic bacteria reveals low species richness and the dominance of *Streptomyces* genus Parthasarathi et al. (2012). Ballav et al. (2015) reported *Streptomyces* spp. as the most predominant group contributing to 46% of the total isolates in crystallizer pond sediments of Ribandar saltern, Goa, India. In this study the isolation was performed with three different types of culture media and four different concentrations of salt because in lagoons were found the concentration of salts between 0-300 psu units (Ballav et al. 2015). Cortes et al. (2019) in the Salar de Huasco report similar results, they were isolated *Streptomyces* (86%), *Nocardiopsis*, (9%), *Micromonospora* (3%), *Bacillu*s (1%) and *Pseudomonas* (1%).

The phylogenetic analysis based on their nearly complete 16S rRNA gene sequences (>1206 bp) was constructed with the isolates M-92, B-81 and B-146 because them presented antibacterial and antiproliferative activities. The phylogenetic tree showed clades well supported to the three isolates and the reference sequences (Figure 3).

**Fig 3.**
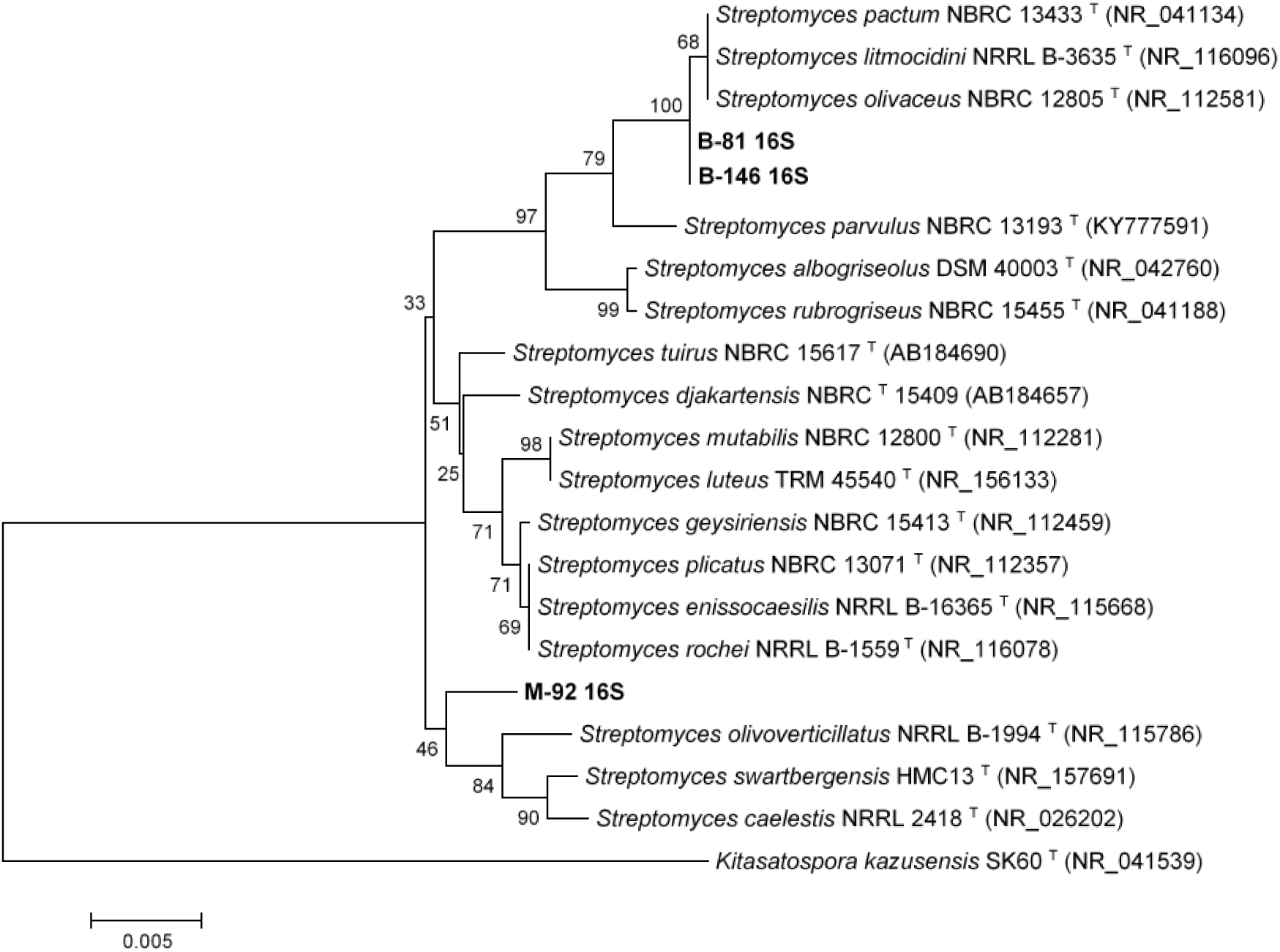
**Phylogenetic tree based on 16S rRNA gene sequences (1206 positions in the final dataset) inferred using the neighbour-joining method in MEGA7, the evolutionary distances were computed using the Kimura 2-parameter method, showing the phylogenetic positions of different saline isolates and type strains within the genus Streptomyces and Pseudonocardia. Numbers at branching points refer to percentages of bootstrap values from 1000 replicates. The scale bar indicates 0.02 indicated substitutions per nucleotide. *Kitasospora kazusensis* SK60^T^ (NR_041539) was used as outgroup.**

Isolate M-92 forms a distinct branch closely related to the type strain of *Streptomyces olivoverticulatus* NRRL-1994^T^, *Streptomyces swartbergensis* HMC13^T^, and *Streptomyces caelestis* NRRL-2418^T^. Isolates B-81 and B-146 form a well-supported sub-clade closely associated to *Streptomyces olivaceus* NBRC 12805^T^, *Streptomyces litmocidini* NRRL B-3635^T^, and *Streptomyces pactum* NBRC 13433^T^ (Figure 3). The length of the branch of all *Streptomyces* isolates in the phylogenetic tree and the assignation to these isolates to completely different clades from each other, highlight the divergence of them from their closely related neighbors. Further studies need to be performed to confirm the right affiliations of these isolates to the novel species within the evolutionary radiation of the genus *Streptomyces*.

### 3.2 Antibacterial activity Screening

Minimum Inhibitory Concentration (MIC) assay was used to screen antibacterial metabolites producer strains against the indicator strains *Escherichia coli* ATCC 10536, *Staphylococcus aureus* ATCC 6538, and *Acinetobacter baumannii* AC-972 MDR; the crude extracts tested from the three extremophilic bacterial reported positive results against the pathogenic bacteria tested. The antimicrobial activity of the crude extract of *Streptomyces* sp. B-81 strain present the highest inhibitory activity with the lowest MIC with 7.82 µg/mL against the three pathogens, followed by *Streptomyces* sp. M-92 showed MIC of 7.82 µg/mL against *E. coli* ATCC 10536 and *S. aureus ATCC 6538*, and 15.63 µg/mL against *A. baumannii AC-972*MDR. *Streptomyces* sp. B-146 showed MIC of 15.63 µg/mL against the pathogens tested. It is important to highlight that all crude extracts of the strains presented low MIC in comparison with the reference antibiotic (Table 4).

**Table 4.** Minimum inhibitory concentration MIC antibacterial of three *Streptomyces* sp. crude extracts, by broth dilution method.

| Isolates | Minimum Inhibition Concentration(ug/mL) |  |  |  |
| --- | --- | --- | --- | --- |
|  | *E. coli*<br>ATCC 10536 | *S. aureus*<br>ATCC 6538 | *A. baumannii*<br>AC-972 MDR | Antibiotic<br>Chloramphenicol |
| M-92 | 7.82 | 7.82 | 15.63 | 22,2 |
| B-146 | 15.63 | 15.63 | 15.63 | 23,1 |
| B-81 | 7.82 | 7.82 | 7.82 | 22,3 |

Similar results were reported in a marine *Streptomyces* sp. S2A with MIC 31.25 µg/mL against *Klebsiella pneumoniae*, 15.62 µg/mL against *Staphylococcus epidermidis, Staphylococcus aureus, Bacillus cereus, Escherichia coli* and *Micrococcus luteus* with 7.8 µg/mL (Saket Siddharth and Ravishankar R. Vittal, 2018). The crude extract of the *Streptomyces* sp. YBQ59 strain against nine pathogens, obtaining MIC results between 10.5 to 22.5 µg/mL, their results being similar to those we obtained (Vu. H et al. 2018).

### 3.3 Antiproliferative activity Screening

The antiproliferative activity of the crude extracts of *Streptomyces* sp. M-92, B-81, and B-146 against three human tumor cell lines were expressed as TGI (Total Growth Inhibition-concentration that inhibited cell growth by 100%) (Table 5).

**Table 5.** Antiproliferative activity of three Streptomyces extracts against human tumor cell lines.

| Crude extract and positive control | TGI (µg/mL)<sup>c</sup> |  |  |
| --- | --- | --- | --- |
|  | U251 | MCF-7 | NCI-H460 |
| Doxorubicin<sup>a</sup> | 2.46 | 6.11 | <0,025 |
| M-92 | 5.06 | 7.96 | 9.33 |
| B-146 | 5.41 | 8.03 | 5.06 |
| Doxorubicin<sup>b</sup> | 1.50 | >25 | 1.80 |
| B-81 | 0.57 | 0.80 | 0.61 |

Human tumor cell lines: U251 (glioma), MCF-7 (breast), NCI-H460 (lung of type non-small cells)
<sup>a, b</sup> Doxorubicin used as positive control;
<sup>c</sup> TGI Concentration that totally inhibited cell growth

According to Abreu et al. (2014), TGI values higher than 50 µg/mL represent inactive samples. In comparison to the others *Streptomyces* tested and doxorrubicin, *Streptomyces* sp. B-81 shown the best result against glioma (TGI =0,57 µg/mL), lung cancer (TGI =0,61 µg/mL) and breast cancer (TGI =0,80 µg/mL). Due to the result obtained with the crude extract of *Streptomyces* sp. B-81 strain, the extracts from three different media were selected for further fractionation, to have access to the metabolic profile. It was possible to verify that the highest activity detected was in the extracts from the fermentation of *Streptomyces* sp. B-81 in the R2A medium (Table 5), which inhibited the nine different types of tumor cell lines. However, it is observed that the fractionation decreases the antitumor activity presented in the crude extract and the putative active compounds are recovered in fractions 3 and 4. Fraction 3 seemed to contain antiproliferative substances in lower concentration and was able to inhibit U251 (glioma) (TGI = 38.3 µg/mL), OVCAR-03 (ovary) (TGI = 62.1 µg/mL), and K562 (leukemia) (TGI = 81.5 µg/mL), and fraction 4 inhibited U251 (glioma) (TGI = 73.5 µg/mL) and UACC-62 (melanoma) (TGI = 89,4 µg/mL) (Table 6).

**Table 6.** **Antiproliferative evaluation of isolate *Streptomyces* sp. B-81 fermented in three different medium and fractions of the R2A medium against nine tumors and one non-tumor cell line.**

| Crude extract | TGI (µg/mL) |  |  |  |  |  |  |  |  |  |
| --- | --- | --- | --- | --- | --- | --- | --- | --- | --- | --- |
|  | 2 | u | m | a | 7 | 4 | o | h | k | q |
| Doxorubicin | 0,03 | 0,76 | >25 | >25 | 0,22 | >25 | 14,4 | >25 | 6,3 | >25 |
| B-81 NB | 70,9 | 250 | * | * | * | * | * | * | * | * |
| B-81 ISP2 | 14,1 | 17,2 | * | 155,9 | * | 38,9 | 55,2 | 155,7 | * | * |
| B-81 R2A | 1,6 | 21,3 | 8,0 | 31,0 | 39,0 | 11,2 | 19,4 | 89,7 | 11,6 | 158,1 |

| B-81 R2A fractions | 2 | u | m | a | 7 | 4 | o | h | k | q |
| --- | --- | --- | --- | --- | --- | --- | --- | --- | --- | --- |
| Fraction 1 | * | * | * | * | * | * | * | * | * | * |
| Fraction 2 | * | * | * | * | * | * | * | * | * | * |
| Fraction 3 | 38,3 | 127,2 | 154,6 | 189,8 | 172,6 | 108,0 | 62,1 | 167,4 | 81,5 | * |
| Fraction 4 | 73,5 | 89,4 | 184,5 | 143,2 | * | 235,1 | 81,6 | 195,2 | 100,4 | * |
| Fraction 5 | * | * | * | * | * | * | * | * | * | * |
| Fraction 6 | 163,0 | * | * | * | * | * | * | * | * | * |

B-81 NB: *Streptomyces* sp. Crude extract in nutrient broth growth medium. B-81 ISP2: *Streptomyces* sp. Crude extract in yeast extract-malt. B-81 R2A: *Streptomyces* sp. Crude extract in R2A growth medium. B-81 R2A fractions: 1 = H<sub>2</sub>O; 2 = MeOH-H<sub>2</sub>O; 3 = MeOH; 4 = MeOH-EtOAc; 5 = 100% EtOAc; 6 = EtOAc -acid: Human tumor lines: 2 = U251 (glioma), u = UACC-62 (melanoma); m = MCF-7 (breast); a = NCI-ADR / RES (ovary with multiple drug resistance phenotype); 7 = 786-O (kidney adenocarcinoma); 4 = NCI-H460 (lung, non-small cell type); o = OVCAR-03 (ovary); h = HT-29 (colon); k = K562 (leukemia). Human non-tumor lineage: q = HaCaT (keratinocyte).
Doxorubicin (Reference chemotherapy).
TGI: Total Growth Inhibition- Concentration necessary to completely inhibit cell proliferation.
\* TGI >250 µg/mL

It is common to find this antiproliferative activity in the *Streptomyces* genus due to its metabolic and genetic capacity to produce secondary metabolites (Tan and Liu, 2017). Additionally, actinomycetes isolated from marine environments have a greater capacity to express these metabolites in response to the pressure of environmental selection to which they have been subjected throughout the evolutionary history of aquatic organisms (Lam, 2006). This pressure has generated high specificity and a complex three-dimensional conformation of the compounds to act in the marine environment (Song et al, 2018). In this particular case, the geographical isolation, the geological conditions of formation of the saline lagoon and the extreme environmental conditions generate a greater selection environmental pressure, which makes it possible to observe antibiotic and antiproliferative activity in the isolates analyzed.

The anticancer potential of halophilic actinomycetes has been extensively studied. Secondary metabolites Salternamides A-D isolated from halophilic actinomycetes of saline on the island of Shinui (Republic of Korea), have potential cytotoxicity against the human colon cancer cell line (HCT116) and gastric cancer cell line (SNU638) (Kim et al. 2015). Another moderately halophilic *Streptomyces sp*. W1 with number accession GenBank JN187420.1, isolated from Weihai Solar Saltern in China, has potent cytotoxicity against human lung cancer cell line (A549) with IC_50_ 78.6 µM; human cervical epithelial cell line (HeLa) IC_50_ 56.6 µM, human liver cancer cell line (BEL-7402) with IC_50_ 47.1 µM, and human colon cancer cell line (HT-29) with IC_50_ 94.3 µM (Liu et al. 2013). Four compounds called shellmicin A-D isolated from the *Streptomyces* sp. shell-016 were described by (Yong Han et al. 2020) and studied concerning cytotoxicity against five human tumor cells lines: non-small cell lung cancer (H1299, ATCC CRL-5803), malignant melanoma (A375, ATCC CRL-1619), hepatocellular carcinoma (HepG2, ATCC HB-8065), colorectal adenocarcinoma (HT29 ATCC HTB-38) and breast cancer (HCC1937, ATCC CRL-2336). The study indicated that the compounds Shellmicin A, B and D showed greater cytotoxicity than Shellmicin C compound, with an IC_50_ ranging from 0.69 µM to 3.11 µM at 72 h. An interesting fact was that shellmicin C and D are a pair of stereoisomers and their biological activity was significantly different. It is worth mentioning that in our study we determined that the activity of the crude extract was better than that of the fractions, the same ones that presented a different activity, therefore, we can deduce that we would be facing compounds with synergism.

An actinomycete strain designated as *Streptomyces* sp. KML-2 isolated from a saline soil mine in Khewra, Pakistan, showed antitumor activity against Hela cells (cervical cancer) and MCF-7 (breast cancer) cell lines with IC_50_ values of 8.9 and 7.8 µg/mL respectively in the initial screening, then they carried out a higher production (20 L) for the purification of the compounds, confirming a wonderful anti-tumor potential against MCF-7 cells with an IC_50_ value of only 0.97 µg /mL (Usman et al. 2015). This study also revealed that the Khewra salt mine from which the KML-2 strain was isolated is a powerful ecological niche with inimitable strain diversity that has yet to be discovered, a fact that reinforces the approach of our study in the search for new compounds with biotechnological activity.

### 3.4 Secondary Metabolite Analysis of *Streptomyces sp*. B-81 isolate

The crude extract obtained from *Streptomyces* sp. B-81 was investigated through UHPLC-MS/MS analyses and demonstrated a broad profile of biomolecules (Fig. S1A). Furthermore, a metabolite screening in the GNPS platform was performed and molecular networking revealed two clusters (A and B) that exhibited secondary metabolites produced by *Streptomyces sp*. B-81 extract (pink) (Fig. 4).

**Fig 4.**
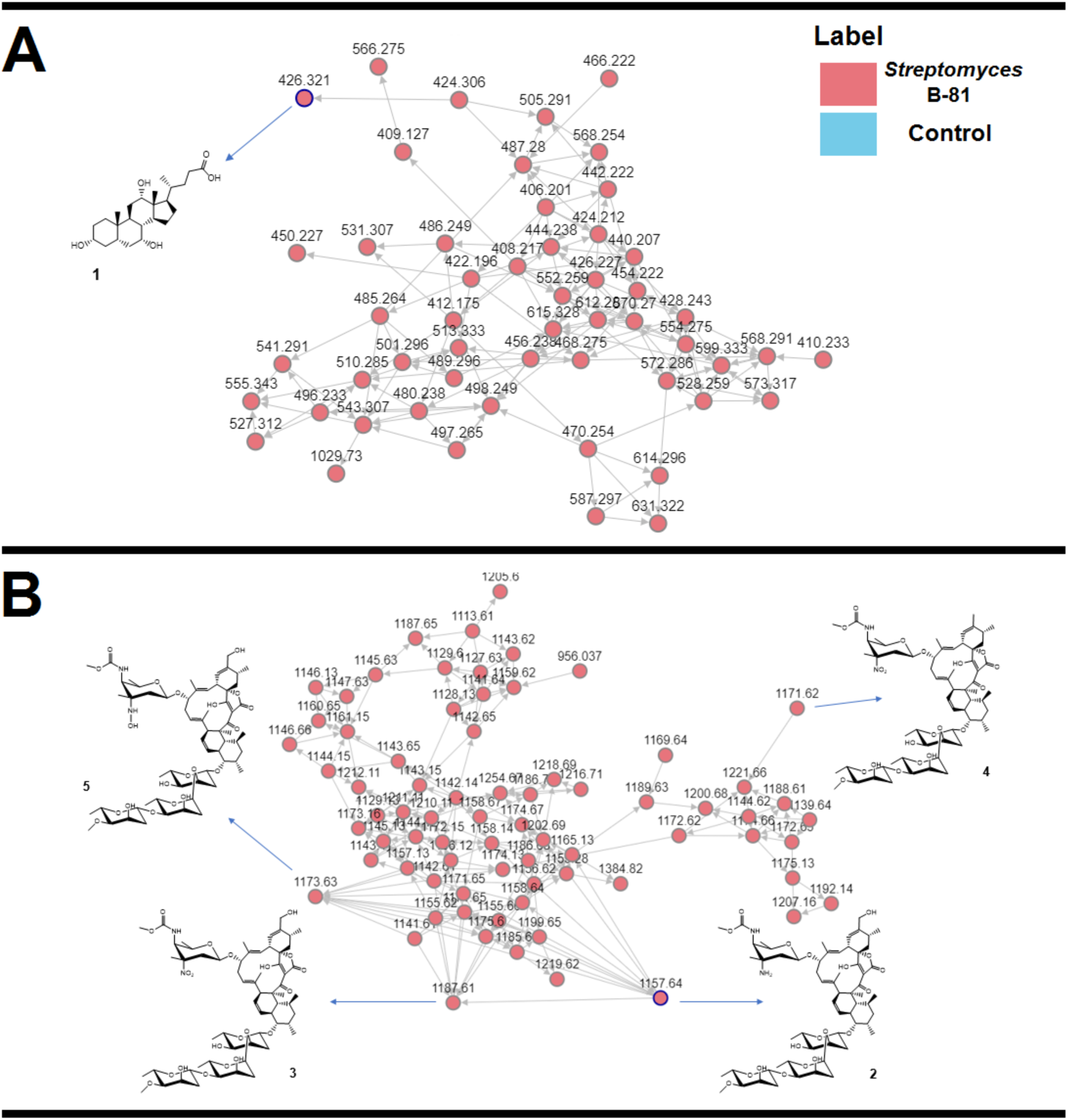
**Molecular networking obtained for *Streptomyces* sp. B-81 extract. Cholic acid and derivatives were grouped in cluster A, and lobophorins were grouped in cluster B. Nodes circled in blue indicate molecules identified by comparison with the GNPS platform database.**

Metabolites were identified as a hit in the GNPS database or manually identified by accurate mass analyses, which showed mass errors below 5 ppm (Table 7).

**Table 7.** **MS data obtained for secondary metabolites detected in *Streptomyces* sp. B-81 extract.**

| Compound | Ion formula | Calculated $m/z$ | Experimental $m/z$ | Error (ppm) |
| --- | --- | --- | --- | --- |
| Cholic acid | C<sub>24</sub>H<sub>44</sub>NO<sub>5</sub> | 426.3219 | 426.3212 | -1.6 |
| Lobophorin A | C<sub>61</sub>H<sub>93</sub>N<sub>2</sub>O<sub>19</sub> | 1157.6372 | 1157.6373 | 0.1 |
| Lobophorin B | C<sub>61</sub>H<sub>91</sub>N<sub>2</sub>O<sub>21</sub> | 1187.6114 | 1187.6108 | -0.5 |
| Lobophorin E | C<sub>61</sub>H<sub>91</sub>N<sub>2</sub>O<sub>20</sub> | 1171.6165 | 1171.6160 | -0.4 |
| Lobophorin K | C<sub>61</sub>H<sub>93</sub>N<sub>2</sub>O<sub>20</sub> | 1173.6322 | 1173.6321 | -0.1 |
| Compound 6 | C<sub>11</sub>H<sub>17</sub>O<sub>3</sub> | 197.1177 | 197.1173 | -2.0 |

The observed signals corresponded to cholic acid (1), lobophorin A (2), lobophorin B (3), lobophorin E (4) and lobophorin K (5).

In cluster A, GNPS database indicated the presence of cholic acid in the *Streptomyces* sp. B-81 extract (Fig. S2 and S3). In the molecular networking, each MS/MS spectra are represented by nodes, which are grouped in clusters based on their fragmentation pattern similarity. Thus, compounds of the same class of molecules are grouped in the same cluster (Wang et al. 2016). As observed in Figure 4, the cholic acid cluster is composed by a range of nodes, suggesting the presence of other analogues. Cholic acid and other bile acids have been reported as secondary metabolites of specific members of the genus *Streptomyces* and *Myroides* (Kim et al. 2007). In the literature, cholic acid-derivatives and bile acids are known as antimicrobials agents, displaying activity against gram-negative and positive bacteria or improving the antimicrobial effect of antibiotics (Schimidt et al. 2001, Rasras et al 2010, Darkoh et al. 2010, Savage et al. 2000).

In cluster B, GNPS database indicated the production of lobophorin A by *Streptomyces* sp. B-81 strain (Fig. S4 and S5). Also, MS/MS profile of compound **2** presented *m/z* 883.49 and 753.43 (Fig. S5), related to the loss of the sugar units in the structure, according with the literature (Nguyen et al. 2020). Similar as observed in cluster A, cluster B also exhibited other lobophorin compounds such as lobophorins B, E and K that were identified based on their accurate masses and fragmentation profiles with typical fragments at *m/z* 184.09, 157.09, 108.08 and 97.06 (Fig. S6-S8). Lobophorins were discovered by Jiang et. al. (1999) in the expedition to Belize on board of the Columbus research ship. In the occasion, a new strain of actinomycete, named as # CNC-837, was isolated from the surface of a brown algae from the Caribbean called *Lobophora variegata*, and was reported to produce lobophorins A and B, potent anti-inflammatory agents (Jiang et al. 1999). Besides lobophorins A and B, other compounds of this class have been reported in the literature. Lobophorins C and D have similar structures to A and B, while Lobophorin C exhibited potent cytotoxic activity against human liver cancer cells, lobophorin D displayed significant inhibitory effect on human breast cancer cells (Wei et al. 2011). Lobophorins E and F were reported by Niu et al. 2011, that also described antibacterial and cytotoxic activities for lobophorin F. In other study, Pan et al. 2013 reported lobophorins H and I and described that lobophorin H exhibited antibacterial activities against *Bacillus subtilis*. Most recent, Lobophorin K was reported (Braña et al. 2017) and also displays antimicrobial and cytotoxic activities.

Besides cholic acid and lobophorins, a furan-type compound (6) were also detected in the *Streptomyces sp*. B-81 extract (Fig. S9). The structures of the metabolites **1**-**6** are shown in Fig. 5.

**Fig 5.**
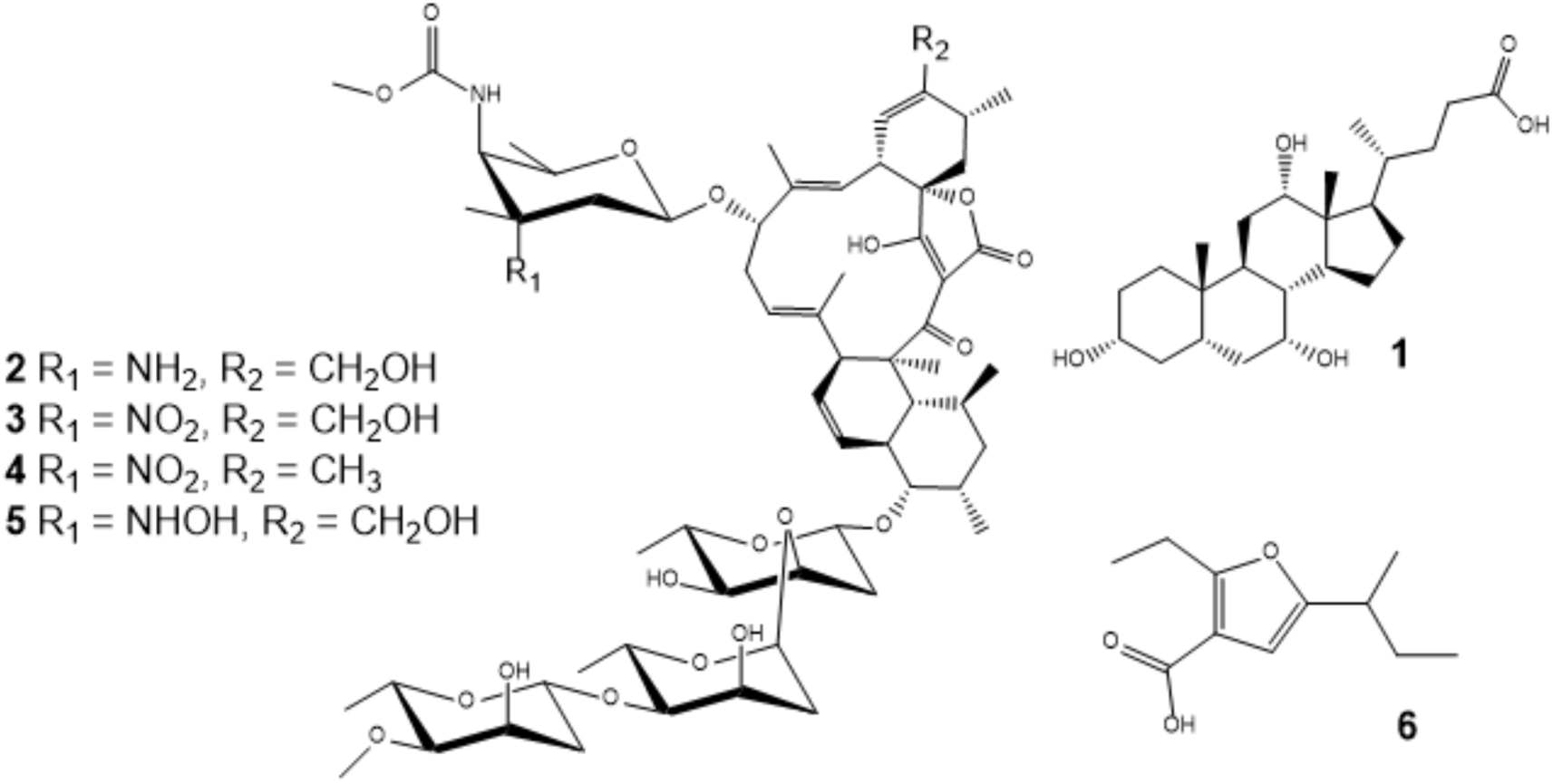
Structures of secondary metabolites identified in *Streptomyces* sp. B-81 extract.

Compound 6 exhibited fragments at m/z 168.07, 167.07 and 153.05, in agreement with the literature (Nguyen et al. 2020). Compound 6 was reported as a secondary metabolite of the *Streptomyces sp*. VN1 strain, a microorganism isolated of sea sediment from a coastal region of Viet Nam (Nguyen et al. 2020). In addition, *Streptomyces sp*. VN1 was also described to produce lobophorin A (Nguyen et al. 2020). Nguyen et al 2020, reported that compound 6 displayed cytotoxic activity, *in vitro*, against 5 types of tumor cell lines. Furthermore, Wang et al. 2008 reported that other furan compound named HS071, produced by *Streptomyces sp*. HS-HY-071, displayed anticancer activity.

Our data deserves further investigation in order to correlate the antimicrobial and cytotoxic activities, observed for the *Streptomyces* sp. B-81 extract, to the bile acids, lobophorins and furan compounds. In addition, the extremophilic environments, in this particular case the Salinas de Bayovar, are prolific sources of microorganisms, such as unique *Streptomyces*. Due the extreme conditions, these microorganisms produce different compounds in order to adapt to survive in the environment, which makes *Streptomyces* sp. B-81 an attractive source of bioactive compounds.

## Conclusion

The present study was successful in determining the diversity and bioactive potential of the actinobacterial isolates present in this type of environment (Mórrope and Bayovar salt flats) located in northern Peru, which are environments that had not been explored for this type of study. this being the first study. Furthermore, it has been found that the isolates of actinobacetria *Streptomyces sp*. M-92, B-146 and B-81 may be new species, their importance for these isolates lies in their antibacterial and antiproliferative potential, being the most promising for their best activity *Streptomyces sp*. B-81. Six biomolecules (colic acid, Lobophorin A, B, E and K in addition to a sixth compound) were detected in this cultivable actinobacterial isolated from Bayovar’s salt flats, but not yet named from the cured base GNPS. With this study it is determined that the actinobacteria that live in these environmental conditions present an enormous wealth of microorganisms that are new species that produce new and biologically active compounds and contribute to the study of extreme environments such as the saline lagoons of northern Peru with antibacterial potential. and antiproliferative. Further studies will be carried out in the identification of these molecules in order to purify and characterize them since this can result in the economic production of bioactive compounds for future pharmaceutical applications.

## Supporting information

Supplemental all

## Acknowledgements

At the research agency of Peru (Concytec-Fondecyt) financially supported this work within the framework of the 041-01 call with the 190-2018 contract; as well as the National Council for Scientific and Technological Development of Brazil (CNPq), it was the authors are grateful to the cooperation of the postgraduate programs in Genetics and Molecular Biology and Biosciences of the Biology Institute of the State University of Campinas. Also, Coordination for the Improvement of Higher Education Personnel - Brazil (CAPES) - (Scholarship-code 001). We thank the research group of the Pluridisciplinary Research Center for Chemistry, Biology and Agriculture to the laboratory infrastructure. Finally, we are also grateful to the anonymous reviewers whose constructive criticisms have significantly improved the quality of the manuscript.

