## Supplemental all for "EVALUATION OF ANTIMICROBIAL AND ANTIPROLIFERATIVE ACTIVITIES OF ACTINOBACTERIA ISOLATED FROM THE SALINE LAGOONS OF NORTHWEST PERU"

**Support Information**

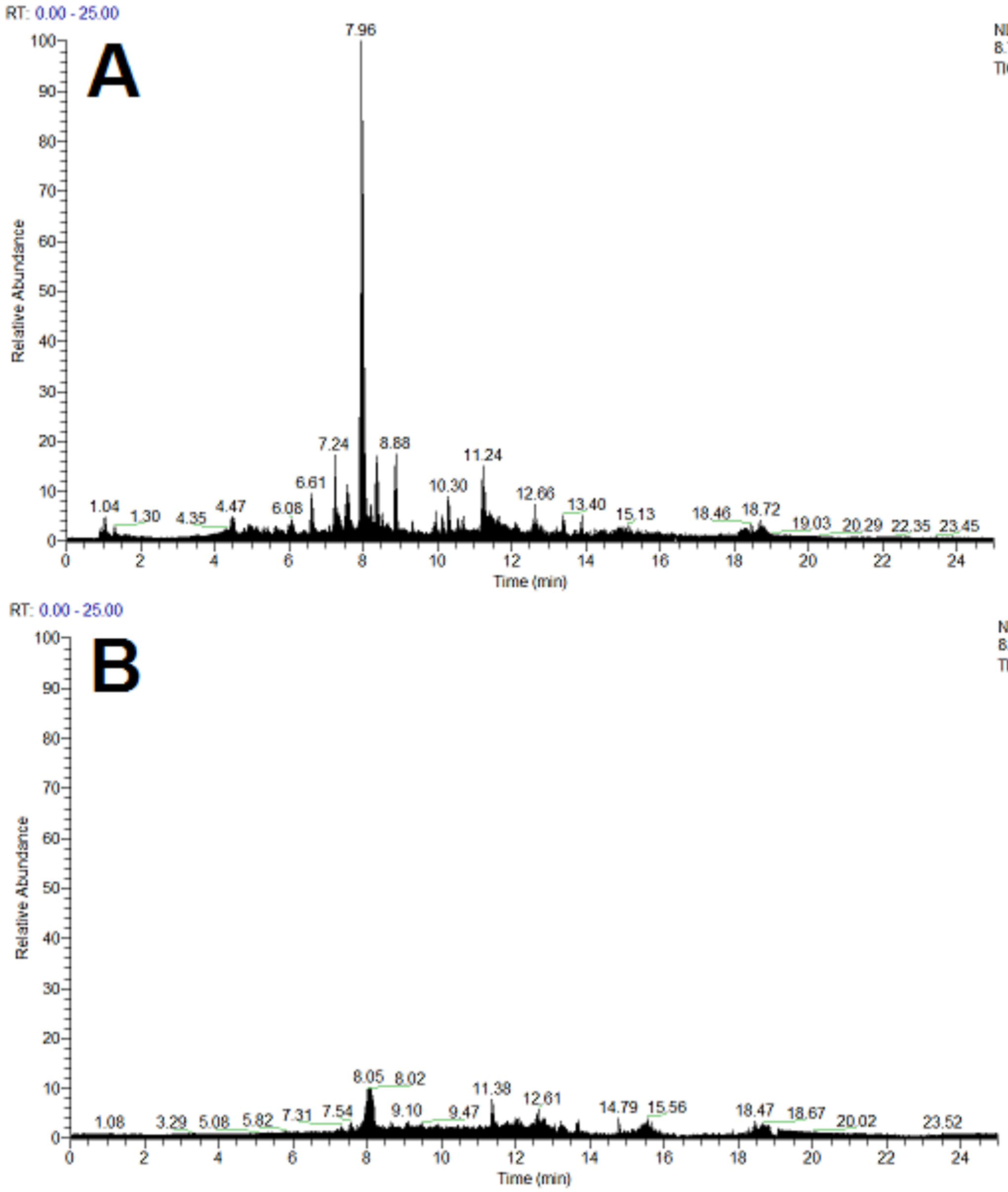


**Fig. S1** Total ion chromatogram (TIC) of UHPLC-MS analyses for (A) *Streptomyces* sp. B-81 extract and (B) control.

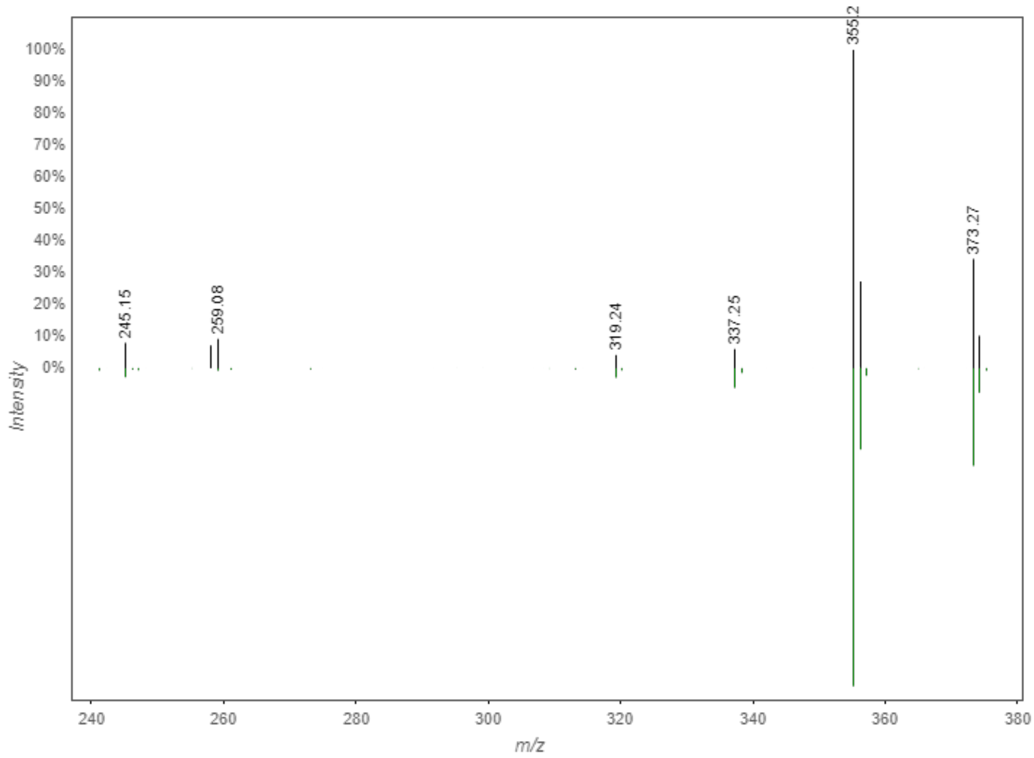


**Fig. S2** MS/MS match between GNPS database (green) and cholic acid (**1**) from *Streptomyces* sp. B-81 extract (black).

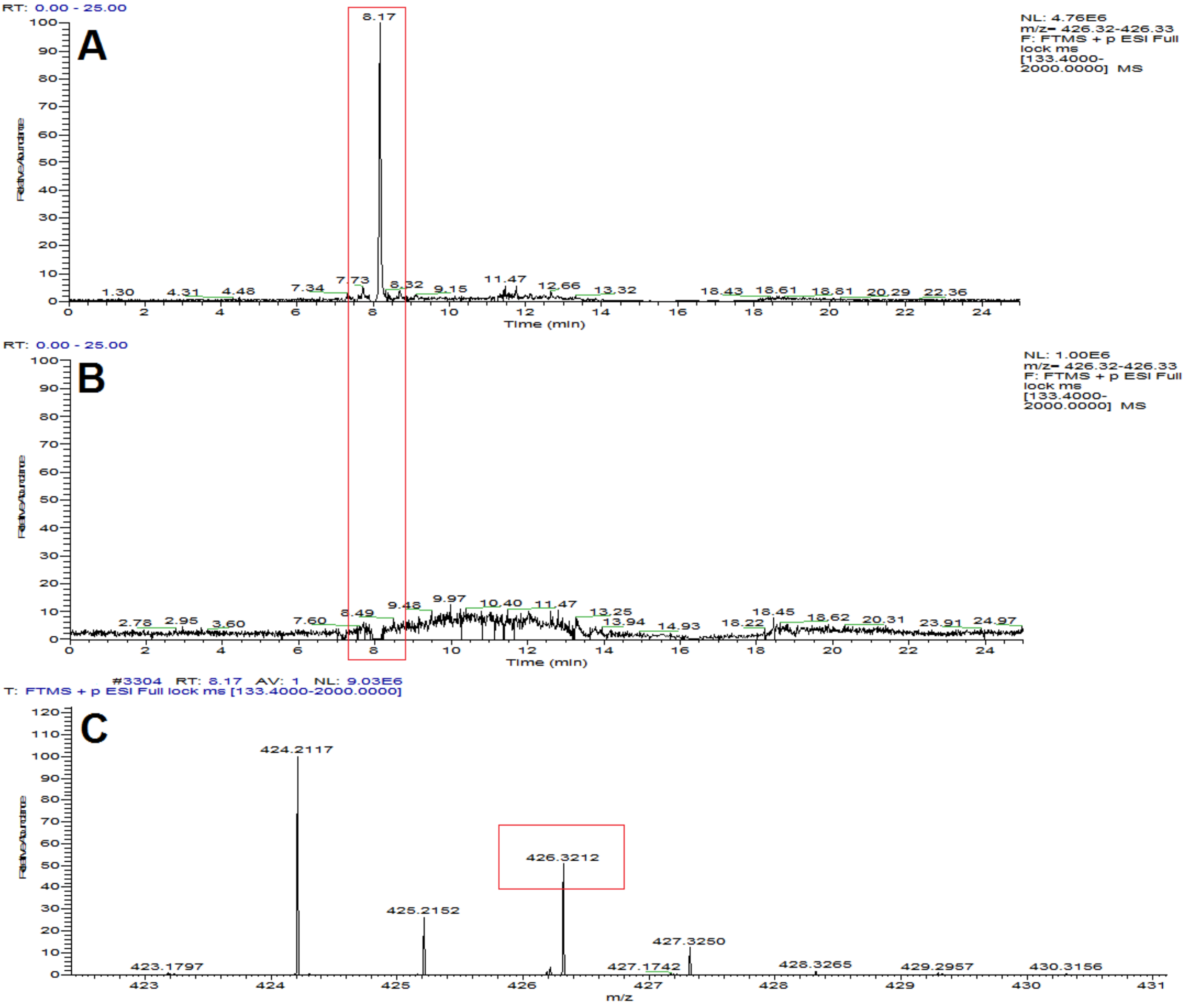


**Fig. S3** Extracted ion chromatograms of *m/z* 426.32 for (A) *Streptomyces* sp. B-81 extract and (B) control. (C) Mass spectrum of ion $[M+NH_4]^+$ *m/z* 426.3212 obtained for compound cholic acid (**1**) (error = -1.6 ppm) at 8.1 min.

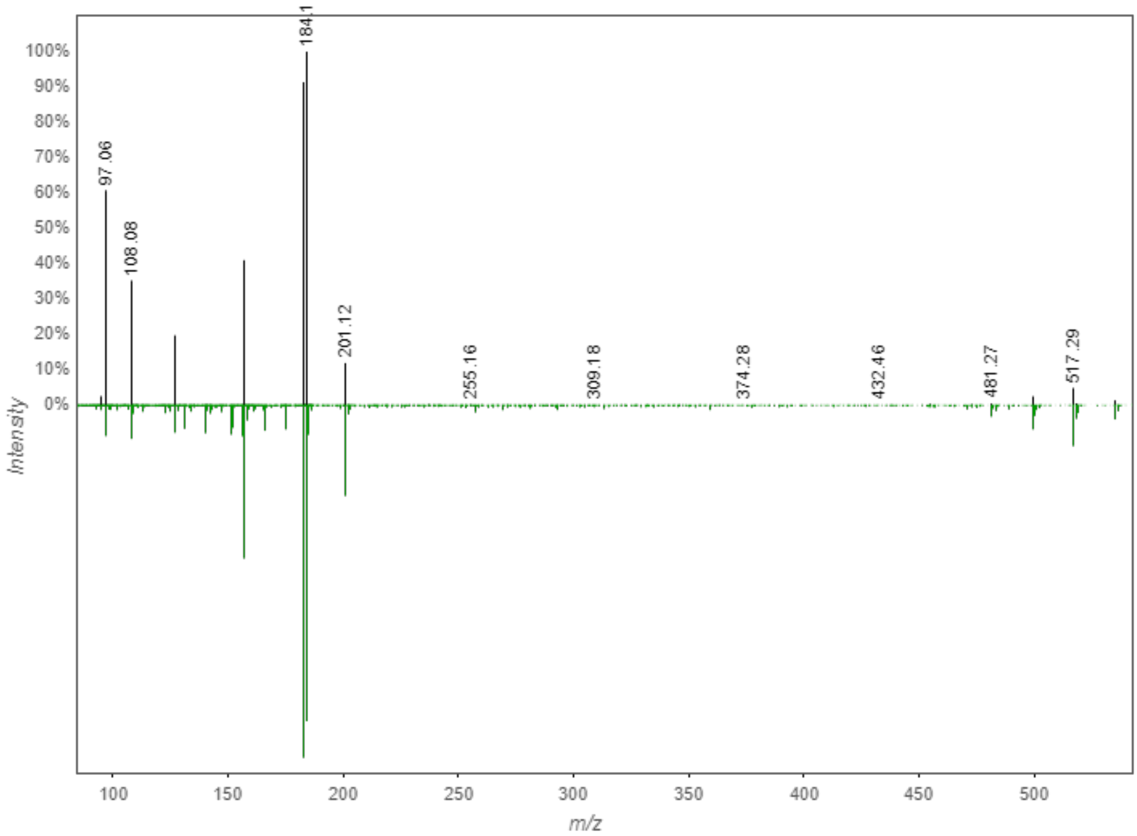


**Fig. S4** MS/MS match between GNPS database (green) and lobophorin A (**2**) from *Streptomyces* sp. B-81 extract (black).

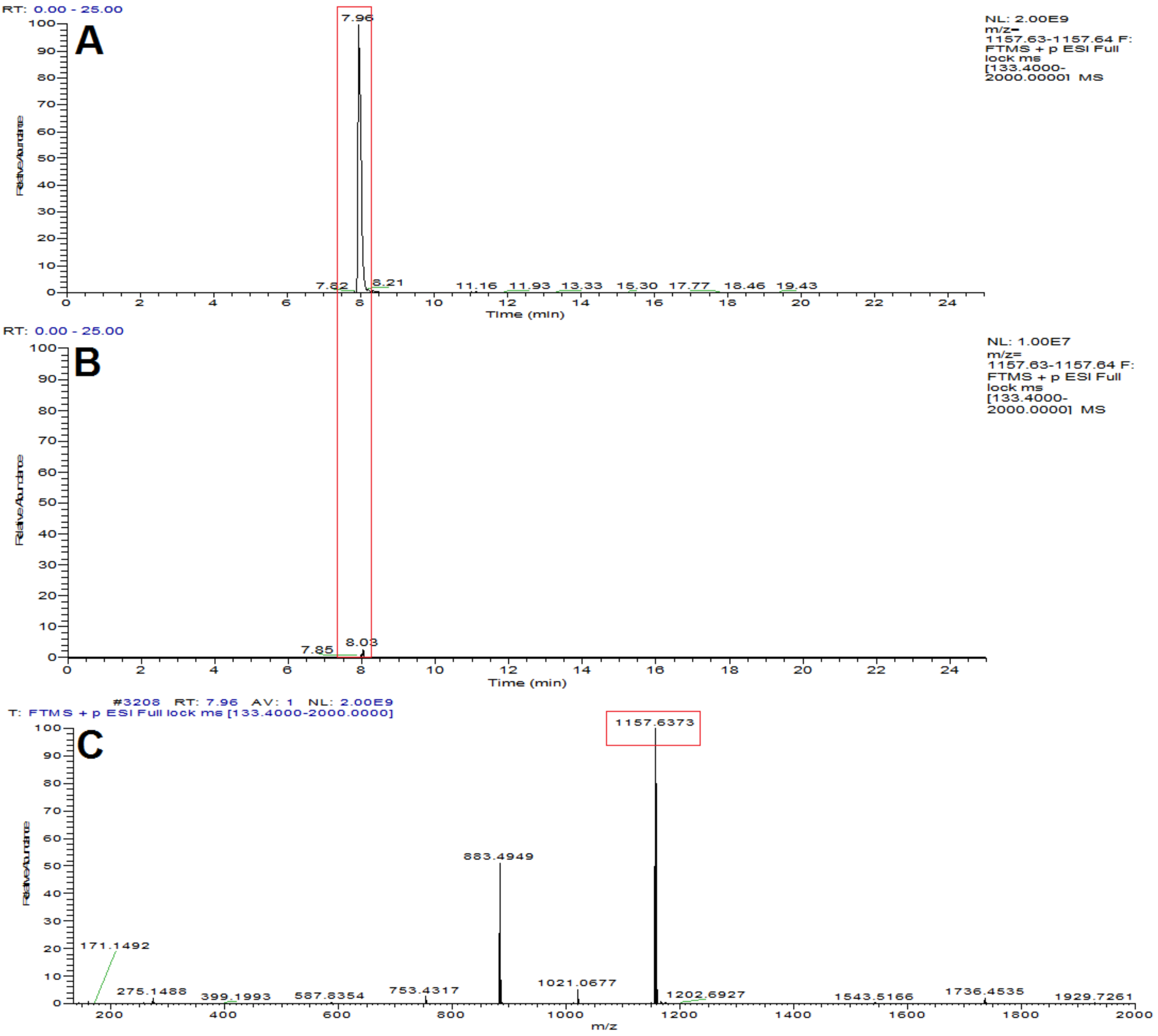


**Fig. S5** Extracted ion chromatograms of *m/z* 1157.63 for (A) *Streptomyces* sp. B-81 extract and (B) control. (C) Mass spectrum of ion $[M+H]^+$ *m/z* 1157.6373 obtained for lobophorin A (**2**) (error = 0.1 ppm) at 7.9 min.

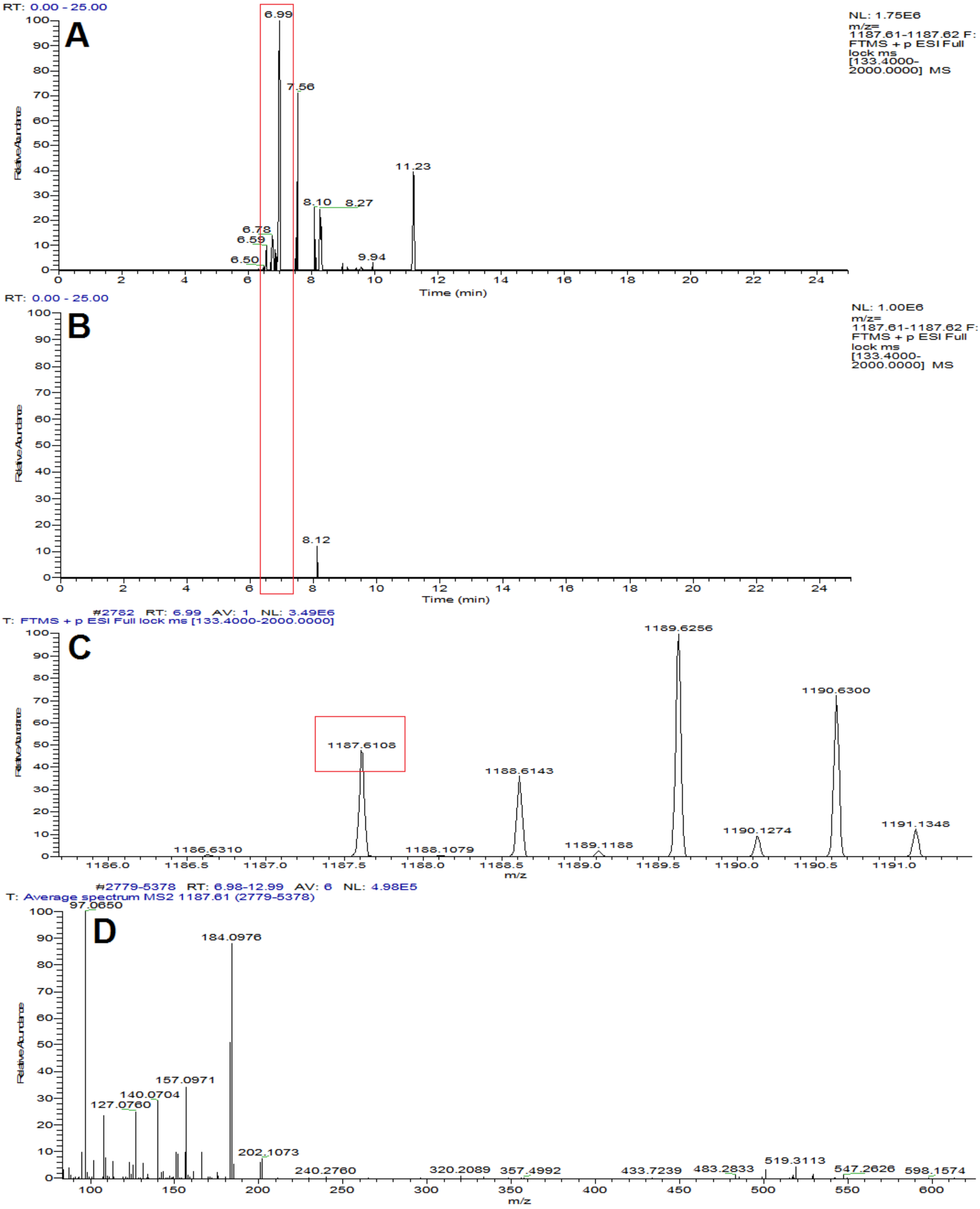

RT: 0.00 - 25.00
A
6.99
7.56
11.23
8.10
8.27
6.78
6.59
6.50
9.94
NL: 1.75E6
m/z= 1187.61-1187.62 F: FTMS + p ESI Full lock ms [133.4000-2000.0000] MS
Relative Abundance
Time (min)
RT: 0.00 - 25.00
B
8.12
NL: 1.00E6
m/z= 1187.61-1187.62 F: FTMS + p ESI Full lock ms [133.4000-2000.0000] MS
Relative Abundance
Time (min)
#2782 RT: 6.99 AV: 1 NL: 3.49E6
T: FTMS + p ESI Full lock ms [133.4000-2000.0000]
C
1189.6256
1190.6300
1187.6108
1188.6143
1190.1274
1191.1348
1189.1188
1186.6310
1188.1079
Relative Abundance
m/z
#2779-5378 RT: 6.98-12.99 AV: 6 NL: 4.98E5
T: Average spectrum MS2 1187.61 (2779-5378)
D
97.0650
184.0976
157.0971
140.0704
127.0760
202.1073
240.2760
320.2089
357.4992
433.7239
483.2833
519.3113
547.2626
598.1574
Relative Abundance
m/z

**Fig. S6** Extracted ion chromatograms of *m/z* 1187.61 for (A) *Streptomyces* sp. B-81 extract and (B) control. (C) Mass spectrum of ion $[M+H]^+$ *m/z* 1187.6108 obtained for lobophorin B (**3**) (error = -0.5 ppm) at 6.9 min. (D) MS/MS spectrum of lobophorin B.

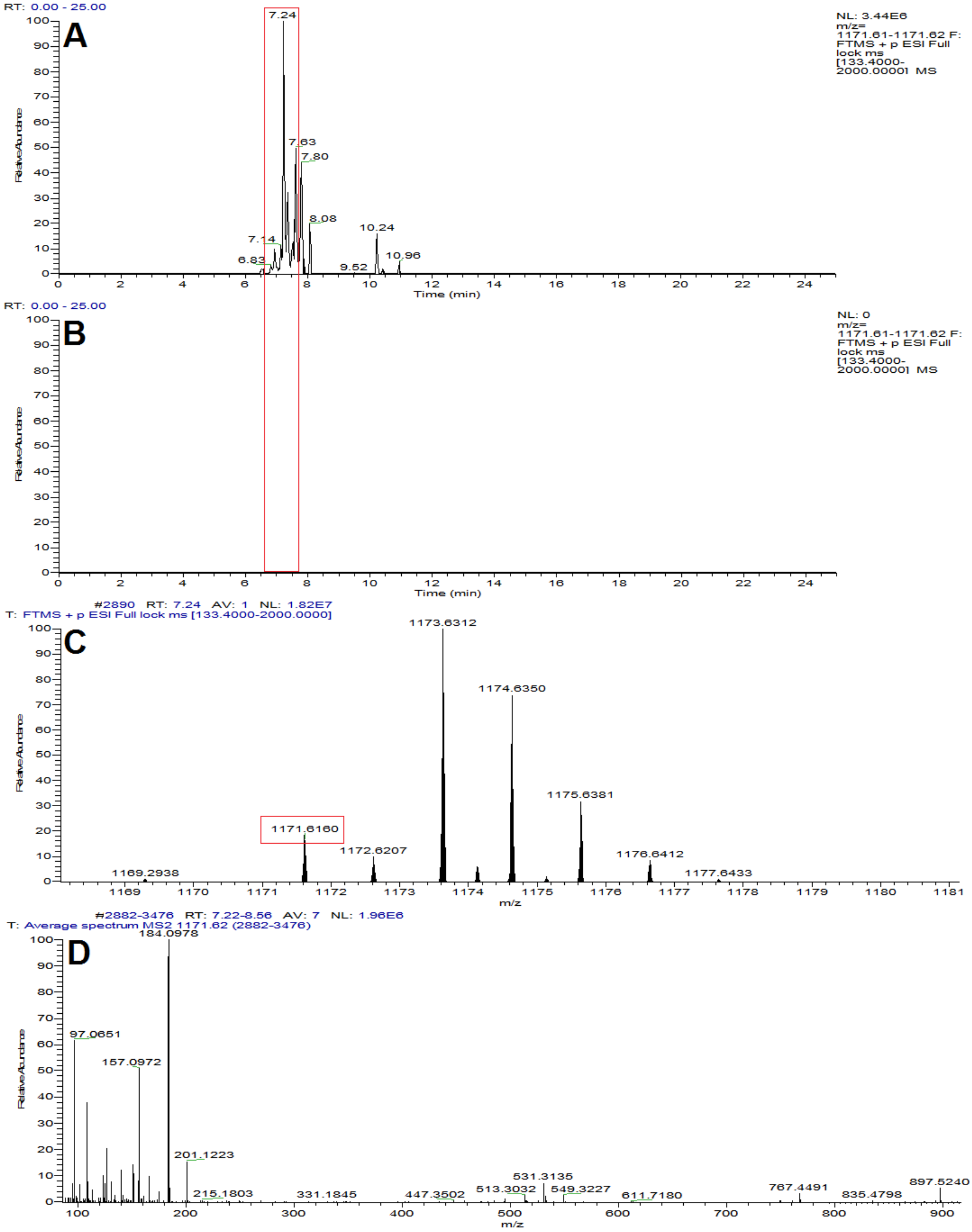

RT: 0.00 - 25.00
A
NL: 3.44E6
m/z= 1171.61-1171.62 F: FTMS + p ESI Full lock ms [133.4000-2000.0000] MS
7.24
7.63
7.80
8.08
10.24
7.14
6.83
9.52
10.96
Relative Abundance
Time (min)
RT: 0.00 - 25.00
B
NL: 0
m/z= 1171.61-1171.62 F: FTMS + p ESI Full lock ms [133.4000-2000.0000] MS
Relative Abundance
Time (min)
#2890 RT: 7.24 AV: 1 NL: 1.82E7
T: FTMS + p ESI Full lock ms [133.4000-2000.0000]
C
1173.6312
1174.6350
1175.6381
1171.6160
1172.6207
1176.6412
1169.2938
1177.6433
Relative Abundance
m/z
#2882-3476 RT: 7.22-8.56 AV: 7 NL: 1.96E6
T: Average spectrum MS2 1171.62 (2882-3476)
D
184.0978
97.0651
157.0972
201.1223
215.1803
331.1845
447.3502
513.3032
531.3135
549.3227
611.7180
767.4491
835.4798
897.5240
Relative Abundance
m/z

**Fig. S7** Extracted ion chromatograms of *m/z* 1171.61 for (A) *Streptomyces* sp. B-81 extract and (B) control. (C) Mass spectrum of ion $[M+H]^+$ *m/z* 1171.6160 obtained for lobophorin E (**4**) (error = -0.4 ppm) at 7.2 min. (D) MS/MS spectrum of lobophorin E.

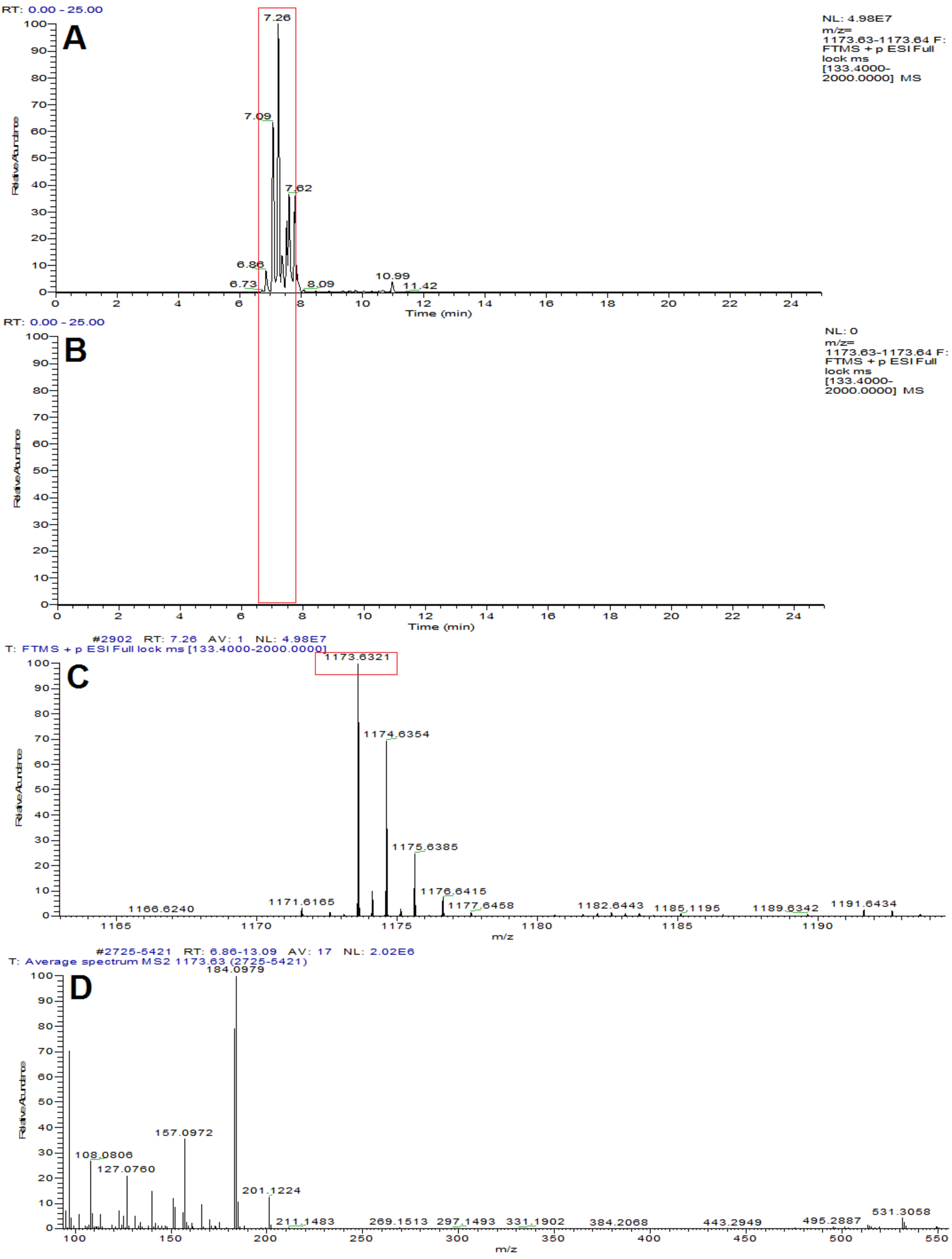

A
B
C
D
RT: 0.00 - 25.00
NL: 4.98E7
m/z= 1173.63-1173.64 F: FTMS + p ESI Full lock ms [133.4000-2000.0000] MS
7.26
7.09
7.62
6.86
6.73
8.09
10.99
11.42
Time (min)
Relative Abundance
NL: 0
#2902 RT: 7.26 AV: 1 NL: 4.98E7
T: FTMS + p ESI Full lock ms [133.4000-2000.0000]
1173.6321
1174.6354
1175.6385
1176.6415
1177.6458
1171.6165
1166.6240
1182.6443
1185.1195
1189.6342
1191.6434
m/z
#2725-5421 RT: 6.86-13.09 AV: 17 NL: 2.02E6
T: Average spectrum MS2 1173.63 (2725-5421)
184.0979
157.0972
108.0806
127.0760
201.1224
211.1483
269.1513
297.1493
331.1902
384.2068
443.2949
495.2887
531.3058

**Fig. S8** Extracted ion chromatograms of *m/z* 1173.63 for (A) *Streptomyces* sp. B-81 extract and (B) control. (C) Mass spectrum of ion $[M+H]^+$ *m/z* 1173.6321 obtained for lobophorin K (**5**) (error = -0.4 ppm) at 7.2 min. (D) MS/MS spectrum of lobophorin K.

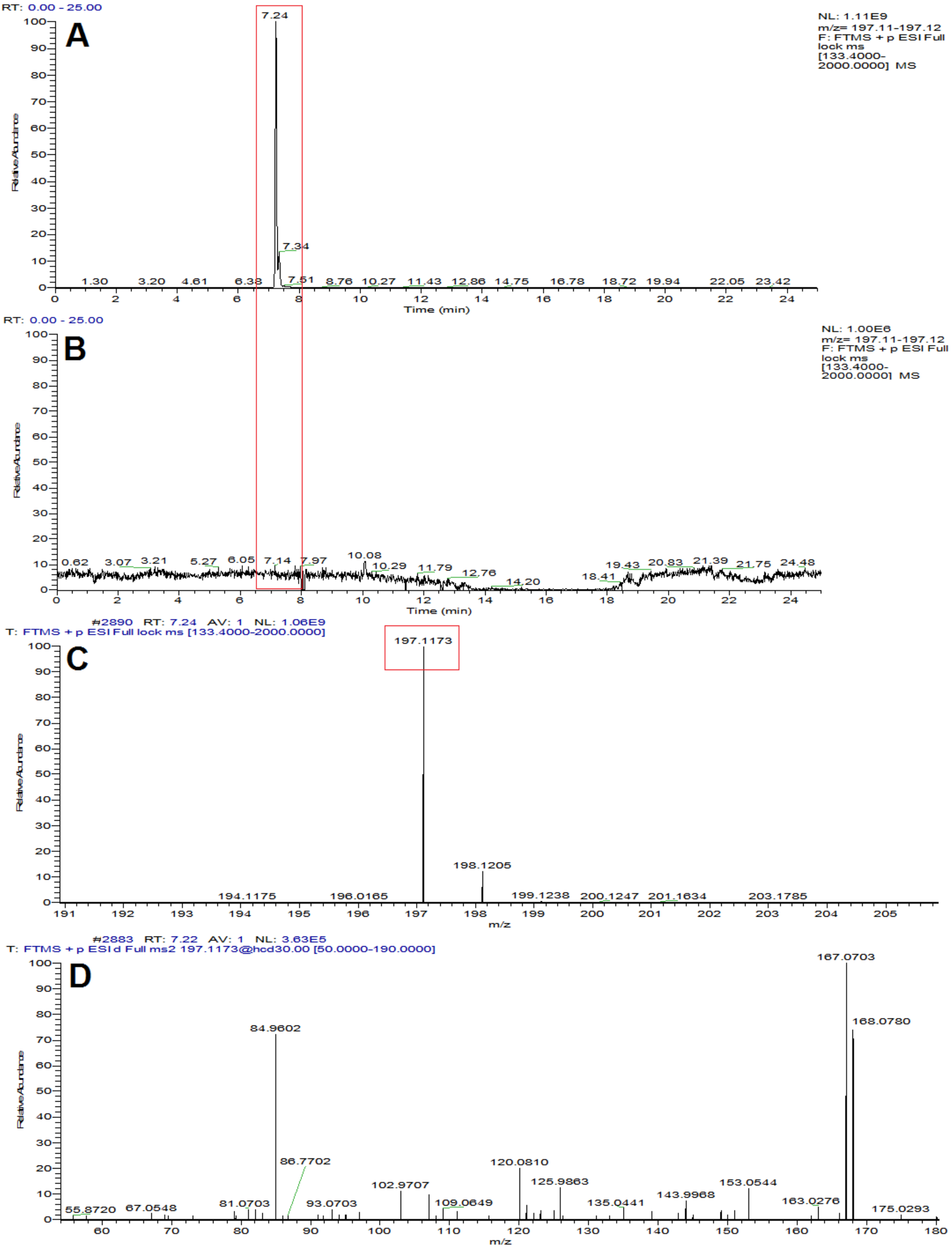

A
RT: 0.00 - 25.00
NL: 1.11E9
m/z= 197.11-197.12
F: FTMS + p ESI Full lock ms [133.4000-2000.0000] MS
Relative Abundance
Time (min)
B
RT: 0.00 - 25.00
NL: 1.00E6
m/z= 197.11-197.12
F: FTMS + p ESI Full lock ms [133.4000-2000.0000] MS
Relative Abundance
Time (min)
C
#2890 RT: 7.24 AV: 1 NL: 1.06E9
T: FTMS + p ESI Full lock ms [133.4000-2000.0000]
197.1173
198.1205
Relative Abundance
m/z
D
#2883 RT: 7.22 AV: 1 NL: 3.63E5
T: FTMS + p ESI d Full ms2 197.1173@hcd30.00 [50.0000-190.0000]
167.0703
168.0780
84.9602
86.7702
120.0810
125.9863
153.0544
102.9707
143.9968
163.0276
Relative Abundance
m/z

**Fig. S9** Extracted ion chromatograms of *m/z* 197.11 for (A) *Streptomyces* sp. B-81 extract and (B) control. (C) Mass spectrum of ion $[M+H]^+$ *m/z* 197.1173 obtained for compound **6** (error = -2.0 ppm) at 7.2 min. (D) MS/MS spectrum of compound **6**.
